# Mutation of lipoprotein processing pathway gene *lspA* or inhibition of LspA activity by globomycin increases MRSA resistance to β-lactam antibiotics

**DOI:** 10.1101/2021.02.03.429649

**Authors:** Claire Fingleton, Merve S. Zeden, Emilio Bueno, Felipe Cava, James P. O’Gara

## Abstract

The *Staphylococcus aureus* cell envelope comprises numerous components, including peptidoglycan (PG), wall teichoic acids (WTA), lipoteichoic acids (LTA), targeted by antimicrobial drugs. MRSA resistance to methicillin is mediated by the *mecA*-encoded β-lactam-resistant transpeptidase, penicillin binding protein 2a (PBP2a). However, PBP2a-dependent β-lactam resistance is also modulated by the activity of pathways involved in the regulation or biosynthesis of PG, WTA or LTA. Here, we report that mutation of the lipoprotein signal peptidase II gene, *lspA*, from the lipoprotein processing pathway, significantly increased β-lactam resistance in MRSA. Mutation of *lgt*, which encodes diacylglycerol transferase (Lgt) responsible for synthesis of the LspA substrate did not impact β-lactam susceptibility. Consistent with previous reports, *lgt* and *lspA* mutations impaired growth in chemically defined media, but not in complex broth. MRSA exposure to the LspA inhibitor globomycin also increased β-lactam resistance. Mutation of *lgt* in an *lspA* background restored β-lactam resistance to wild type. The *lspA* mutation had no effect on PBP2a expression, PG composition or autolytic activity indicating a potential role for WTA or LTA. The *lspA* and *lgt* mutants exhibited marginally increased resistance to the D-alanine pathway inhibitor D-cycloserine. In addition, mutation of *lgt* and multicopy *lspA* expression, but not mutation of *lspA*, significantly increased susceptibility to the lipoteichoic acid synthase inhibitor Congo red revealing complex interplay between lipoprotein processing mutations and the expression/stability of cell surface glycopolymers. These findings indicate that accumulation of the LspA substrate, diacylglyceryl lipoprotein, increases MRSA resistance to β-lactam antibiotics through impacts on cell envelope components other than PG.

## Introduction

The cell envelope of *Staphylococcus aureus* comprises a cytoplasmic membrane surrounded by a thick peptidoglycan layer, cell wall-anchored proteins, lipoteichoic acids (LTA), wall teichoic acids (WTA) and cell surface proteins. Accurate biosynthesis, assembly and stability of these cell envelope components is essential for the growth and pathogenesis of *S. aureus*, and is the target of numerous antimicrobial agents (1). The peptidoglycan layer determines cell shape and protects the cell from osmotic lysis, cell surface proteins have important roles in adhesion, biofilm formation, and immune evasion, and teichoic acids are involved in protecting the cell from the activity of cationic antimicrobial peptides.

Methicillin resistance in MRSA is mediated by the *mecA*-encoded, low-affinity penicillin-binding protein 2a (PBP2a) carried on the mobile staphylococcal cassette chromosome *mec* resistance (HeR) under laboratory growth conditions. HeR strains can become highly or homogeneously resistant (HoR) after selection on elevated β-lactam concentrations via poorly understood mechanisms, which require accessory mutations at other chromosomal loci frequently associated with activation of the stringent response, cyclic-di-adenosine monophosphate (c-di-AMP) signalling pathway (2-5), the activity of RNA polymerase (6) and the ClpXP chaperone-protease complex (7,8). In addition, methicillin susceptible *S. aureus* strains (MSSA) lacking *mecA* can also acquire low-level resistance through adaptive mutations impacting the c-di-AMP signalling pathway and ClpXP activity (9).

Bacterial lipoproteins are a class of lipid-modified membrane proteins, involved in a range of diverse functions such as; nutrient acquisition (10), signal transduction (11), respiration (12), protein folding (13), virulence (14), antibiotic resistance (15) and host invasion (16). Mature lipoproteins are composed of lipid moieties, specifically acyl groups, linked to the N-terminus of a protein. The hydrophobic nature of the acyl groups serves as a membrane anchor for the lipoprotein (17). In Gram-negative bacteria, lipoproteins reside in both the cytoplasmic and outer membranes, while in Gram-positive bacteria they are anchored in the outer leaflet of the cytoplasmic membrane and the protein portion may extend into the cell wall and beyond (18).

Lipoprotein genes are estimated to comprise 1-3% of all genes in bacterial genomes (18). While many lipoproteins have been identified and experimentally validated, others are putatively identified using predictive software, thus their functions remain unknown (19). A recent bioinformatic evaluation of Staphylococcal lipoproteins in the MRSA strain USA300 identified 67 lipoproteins, comprising 2.57% of all genes (18). When grouped by function, 25 of the 67 lipoproteins were implicated in ion (notably iron) and nutrient transport, 8 were ascribed miscellaneous functions including sex pheromone biosynthesis, respiration, chaperone-folding, and protein translocation and 15 were classified as tandem lipoproteins, of which 9 are “lipoprotein-like” lipoproteins, known to play a role in host cell invasion (16). The remaining 19 lipoproteins were not assigned any known function.

Lipoproteins are synthesised in a precursor form called preprolipoproteins. The N-terminal domain includes a type II signal peptide, approximately 20 amino acids in length (20), which enables translocation of preprolipoproteins to the cytoplasmic membrane, predominantly via the general secretory (Sec) pathway (21). The signal peptide has 3 distinct domains; a positively charged N domain, a hydrophobic H domain and a C-terminal lipobox. The lipobox is comprised of a conserved 3-amino acid sequence [LVI]_-3_ [ASTVI]_-2_ [GAS]_-1_ in front of an invariant cysteine residue [C]_+1_ (22,23). This lipobox serves as a recognition site for enzymes of the lipoprotein processing pathway, enabling lipid modification of the cysteine residue, and cleavage of the signal peptide between the amino acid at position −1 and the +1 cysteine (22). The first enzyme in the lipoprotein processing pathway is diacylglycerol transferase (Lgt) which covalently attaches a diacylglycerol molecule from phosphatidyl glycerol onto the sulfhydryl group of the invariant cysteine, resulting in a prolipoprotein (24). This diacylglycerol serves as a membrane anchor. Next, the type II lipoprotein signal peptidase (Lsp) cleaves the signal peptide between the amino acid at position −1 and +1, leaving the invariant cysteine residue as the new terminal amino acid (25). Lgt and Lsp are conserved in all bacterial species. In Gram-negative bacteria, a third step is catalysed by the enzyme N-acyl transferase (Lnt), which transfers an N-acyl group onto the invariant cysteine residue at the N-terminal of the protein (26). Lnt homologs have been identified in high-GC Gram-positive bacteria (27) but not in low-GC Firmicutes. Despite the lack of an apparent Lnt homolog, N-acylated lipoproteins have been identified in *S. aureus* (28) and recent work has identified two novel non-contiguous genes *InsA* and *InsB* which catalyse the N-terminal acylation of lipoproteins in *S. aureus* (29). The membrane metalloprotease Eep and the EcsAB transporter were shown to be involved in the processing and export of linear peptides, including the signal peptide cleaved by LspA in the lipoprotein processing pathway (30–32). Lgt and Lsp are essential for the viability of Gram-negative bacteria (33). In contrast, *lgt* and *lspA* mutations do not impact viability in Gram-positive bacteria (34), but are associated with changes in growth, immunogenicity (35) and virulence (14) phenotypes.

In this study, we characterised the impact of *lspA and lgt* mutations, alone and in combination, on susceptibility to β-lactams, D-cycloserine and Congo red, growth, PBP2a expression, peptidoglycan structure, and autolytic activity in MRSA. The impact of globomycin, which is known to inhibit LspA activity, on β-lactam susceptibility was also characterised. Our data suggest that accumulation of the LspA substrate, diacylglyceryl-prolipoprotein, modulates resistance to β-lactam antibiotics in MRSA.

## Results

### Mutation of *lspA* in MRSA increases resistance to β-lactam antibiotics

The Nebraska Transposon Mutant Library (NTML) (36) was screened to identify mutants exhibiting altered susceptibility to cefoxitin, which is recommended as a surrogate for measuring *mecA-* mediated oxacillin resistance in clinical laboratories, in accordance with Clinical and Laboratory Standards Institute (CLSI) guidelines for disk diffusion susceptibility assays. Mutants identified by this screen included NE869 *(yjbH)* (37), NE1909 *(sagA)* (38) and NE810 (*cycA*)(39), all of which have previously been implicated in β-lactam resistance. A new mutant identified in this screen was NE1757 (*lspA*::Tn), which exhibited increased resistance to cefoxitin (Fig 1A, B). PCR was used to confirm the presence of *lspA*::Tn allele in NE1757 (data not shown). Using E-test strips, as described in the methods, oxacillin MIC of the *lspA* mutant NE1757 was found to be 128 - 256 μg/ml, compared to 32 - 64 μg/ml for JE2 (Fig. 1C). Two JE2 transductants carrying the *lspA*::Tn allele from NE1757 also exhibited increased resistance to oxacillin (Fig. 1C).

**Fig. 1.**
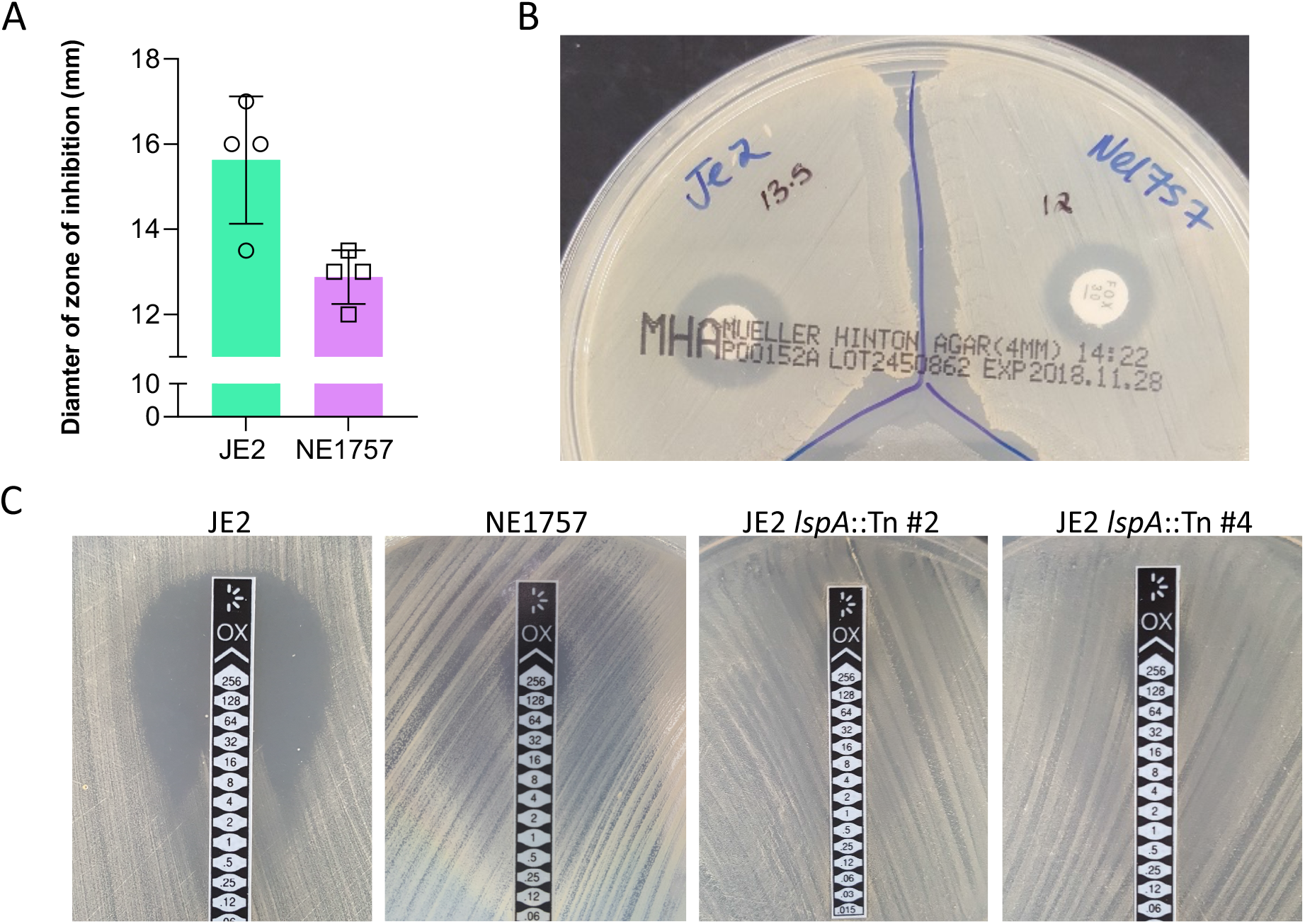
Mutation of *lspA* increases resistance to cefoxitin and oxacillin. **(A)** Average diameters of the cefoxitin disk zones of inhibition for JE2 and NE1757 (*lspA*::Tn) from 4 independent experiments, plotted using Prism software (GraphPad). **(B)** Representative image of JE2 (left) and NE1757 (right) grown on MH agar with a cefoxitin 30 μg disk. **(C)** M.I.C.Evaluator measurement of oxacillin minimum inhibitory concentrations (MICs) in JE2, NE1757 (*lspA*::Tn), and two independent JE2 transductants (#2 and #4) carrying the *lspA*::Tn allele grown on MHB 2% NaCl agar. This assay was repeated 3 independent times for each strain and a representative image is shown.

The increased oxacillin resistance phenotype of NE1757 was also complemented by the introduction of a plasmid (pLI50)-borne copy of the wild type *lspA* gene *(plspA)* into the mutant. Growth of JE2, NE1757, NE1757 pLI50 and NE1757 *plspA* on MHA 2% NaCl supplemented with oxacillin 32 μg/ml visually demonstrated that carriage of the complementation plasmid reversed the increased oxacillin resistance phenotype of NE1757 (Fig. S1). Measurement of oxacillin MICs by agar dilution showed that NE1757 and NE1757 pLI50 had MICs of 256 μg/ml, while JE2 and the complemented strain NE1757 *plspA* had MICs of 64 μg/ml (Table 1).

**Table 1.**
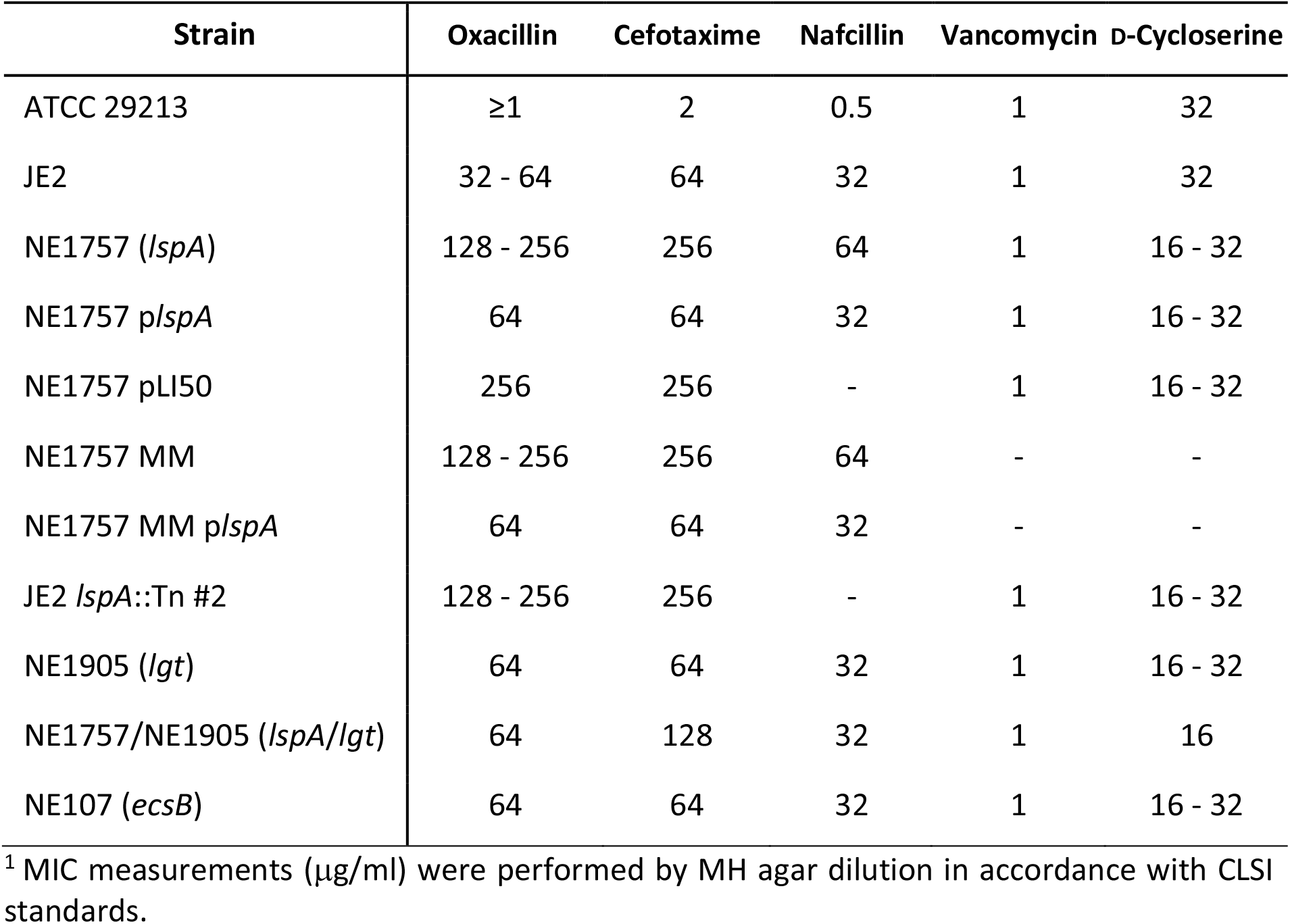
Antibacterial activity of oxacillin, cefotaxime, nafcillin, vancomycin and D-cycloserine (MIC measurements; μg/ml^1^) against ATCC 29213 (control), JE2, NE1757 (*lspA*::Tn), NE1757 *plspA*, NE1757 pLI50, NE1757 MM (markerless *lspA* mutant), NE1757 MM *plspA*, JE2 *lspA*::Tn #2 (transductant), NE1905 (*lgt*::Tn), NE1757/NE1905 double mutant *lspA/lgt* and NE107 (*ecsB*).

Comparison of JE2 and NE1757 growth in MHB, MHB 2% NaCl, TSB and TSB 0.5 mg/ml oxacillin revealed no significant differences (Fig. S2). Similarly, population analysis profiling revealed that the heterogeneous pattern of oxacillin resistance expressed by JE2 was unchanged in NE1757 (Fig. S3). These observations indicate that the increased β-lactam resistance phenotype of NE1757 was not attributable to any growth advantage or change in the heterogeneous/homogeneous oxacillin resistance profile.

Comparative WGS analysis confirmed that the only change in the NE1757 genome was the insertion of the *Bursa aurealis* transposon in the *lspA* gene at position 1192002 (Table S1) and there were no SNPs present. The NE1757 genome was also checked manually for zero coverage regions to confirm the absence of any large deletions and insertions. Taken together these data indicate that mutation of *lspA*, which encodes lipoprotein signal peptidase II involved in the lipoprotein processing pathway (Fig. 2), increases resistance to oxacillin and cefoxitin in JE2.

**Fig. 2.**
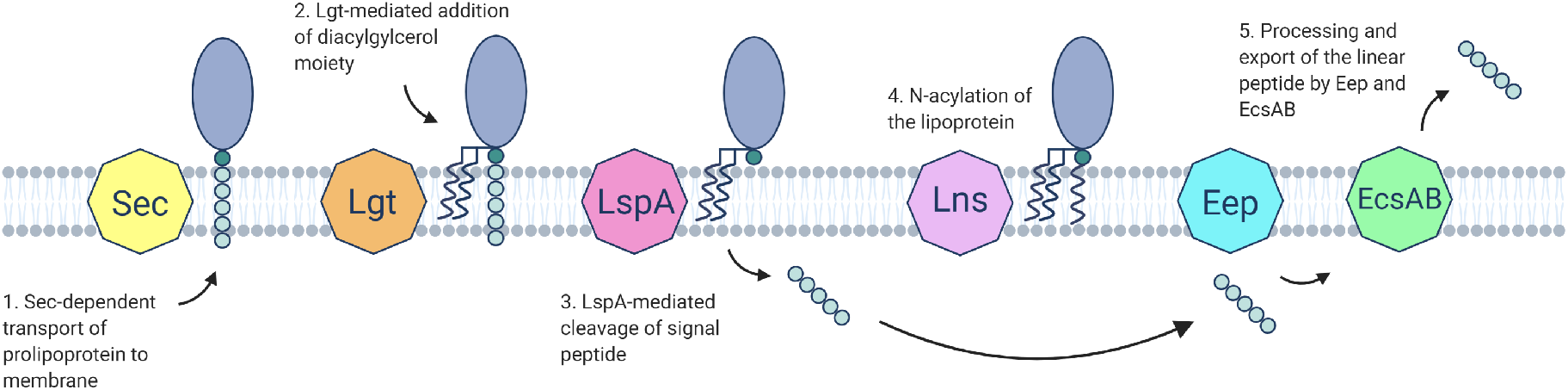
Role of LspA in the proposed model of lipoprotein processing in Gram-positive bacteria. Lipoproteins are synthesized in the cytoplasm as precursors. **1.** Prolipoprotein is translocated usually via the Sec machinery through recognition of its signal peptide (SP). This sequence also contains the lipobox specific for bacterial lipoproteins. **2.** Diacylglyceryl transferase (Lgt) catalyzes the transfer of a diacylglyceryl group from phosphatidylglycerol onto the prolipoprotein resulting in diacylglyceryl-prolipoprotein. **3.** Lipoprotein signal peptidase II (LspA) recognizes the diacylglyceryl-modified signal peptide and cleaves between the amino acid at position-1 and the lipid-modified cysteine residue at +1. **4.** The lipoprotein N-acylation transferase system (Lns) converts diacyl lipoproteins to triacylated lipoprotein (29). **5.** Eep and EcsAB play roles in the processing and secretion of linear peptides (32).

### Mutation of *lspA* does not affect PBP2a expression or peptidoglycan structure and crosslinking

One of the numerous lipoproteins processed by LspA is PrsA, a chaperone and foldase protein, reported to play a role in PBP2a folding and β-lactam resistance (40). To investigate if improper processing of PrsA, or another lipoprotein, in the *lspA* mutant impacted on PBP2a, Western blotting was used to compare PBP2a expression in JE2, NE1757 and NE1757 *plspA.* The MSSA strain 8325-4 was included as a *mecA*-negative control. Growth of JE2 and NE1757 in MHB without 2% NaCl at 37°C (data not shown) or MHB with 2% NaCl at 35°C supplemented with 0.5 μg/ml oxacillin (Fig. 3A) revealed similar levels of PBP2a expression in all strains.

**Fig. 3.**
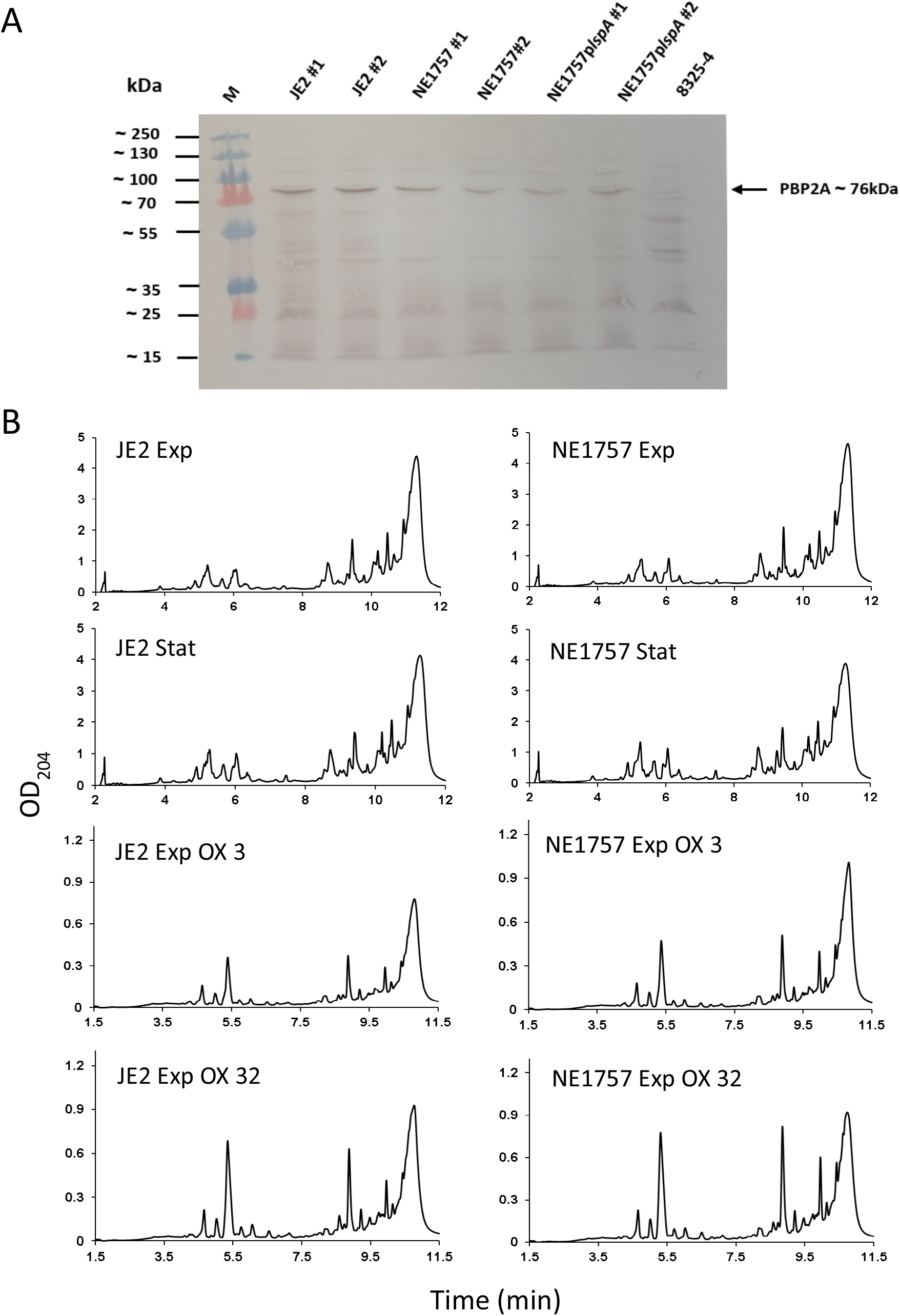
Mutation of *lspA* does not affect PBP2a expression levels or peptidoglycan structure and crosslinking. **(A)** Western blot of PBP2a protein in JE2, NE1757 (*lspA*), NE1757 *plspA* and MSSA strain 8325-4 (negative control). Cells were grown to exponential stage in MHB 2% NaCl supplemented with 0.5 μg/ml oxacillin, with the exception of 8325-4 which was grown in MHB 2% NaCl. For each sample, 8 μg total protein was run on a 7.5% Tris-Glycine gel, transferred to a PVDF membrane and probed with anti-PBP2a (1:1000), followed by HRP-conjugated protein G (1:2000) and colorimetric detection with Opti-4CN Substrate kit. Three independent experiments were performed and a representative image is shown. (**B)** Representative UV chromatograms of peptidoglycan extracted from JE2 and NE1757 collected from cultures grown to exponential or stationary phase in MHB or MHB supplemented with oxacillin 3 μg/ml or 32 μg/ml. Each profile shown is a representative of 3 biological replicates.

Quantitative peptidoglycan compositional analysis was performed using UPLC analysis of muramidase-digested muropeptide fragments extracted from exponential or stationary phase cultures of JE2 and NE1757 grown in MHB or MHB supplemented with oxacillin 3 μg/ml or 32 μg/ml. The PG profile of JE2 and the *lspA* mutant NE1757 were similar under all growth conditions tested (Fig. 3B). Thus, supplementation of MHB with oxacillin was associated with significant changes in muropeptide oligomerization and reduced crosslinking, but these effects were the same in both JE2 and NE1757 (Fig. S4). The total PG concentrations extracted from JE2 and NE1757 cell pellets were also the same (data not shown). Comparison of Triton X-100-induced autolysis in JE2 and NE1757 also revealed identical autolytic profiles (Fig. S5). Finally, the NaCl tolerance phenotypes of JE2 and NE1757 were also similar (Fig. S6), indicating that c-di-AMP signalling, which has previously been implicated in the control of β-lactam resistance, autolytic activity and NaCl tolerance (4,5) was unaffected by the *lspA* mutation. These data indicate that increased β-lactam resistance in the *lspA* mutant was not associated with significant changes in PG abundance, structure, crosslinking, c-di-AMP signalling or autolytic activity.

### Exposure to the LspA inhibitor globomycin also increases β-lactam resistance

Globomycin is a natural peptide antibiotic, first discovered in 1978, produced by 4 different strains of actinomycetes (41,42). It is an inhibitor of LspA and works by sterically blocking the active site of the enzyme (43). Globomycin has moderate to strong antibacterial activity against many Gram-negative species and has been proposed to cause disruption of cell surface integrity (44). However, despite its ability to inhibit LspA, globomycin does not have significant antimicrobial activity against Gram-positive bacteria including *S. aureus*, with MICs >100 μg/ml (41,42,45).

Because disruption of *lspA* increased resistance to β-lactams, we hypothesized that chemical inhibition of LspA by globomycin may also be associated with increased β-lactam resistance. To test this hypothesis, the susceptibility of JE2 and NE1757 to oxacillin was determined in the presence or absence of globomycin. A series of JE2 and NE1757 cultures grown in MHB 2% NaCl were used to determine that oxacillin 40 μg/ml inhibited growth of JE2 but not NE1757 (Fig. 4A) as predicted. Next, JE2 MHB 2% NaCl 40 μg/ml oxacillin cultures were further supplemented with 10, 20, 30, 40 or 50 μg/ml globomycin (Fig. 4B). Oxacillin-induced inhibition of JE2 growth was rescued globomycin, optimally at 10, 20 and 30 μg/ml (Fig. 4B). Growth of JE2 in 40 μg/ml oxacillin and 50 μg/ml globomycin was substantially impacted compared to lower globomycin concentration (Fig. 4B) indicating that the antagonism of oxacillin by globomycin was dose dependent.

**Fig. 4.**
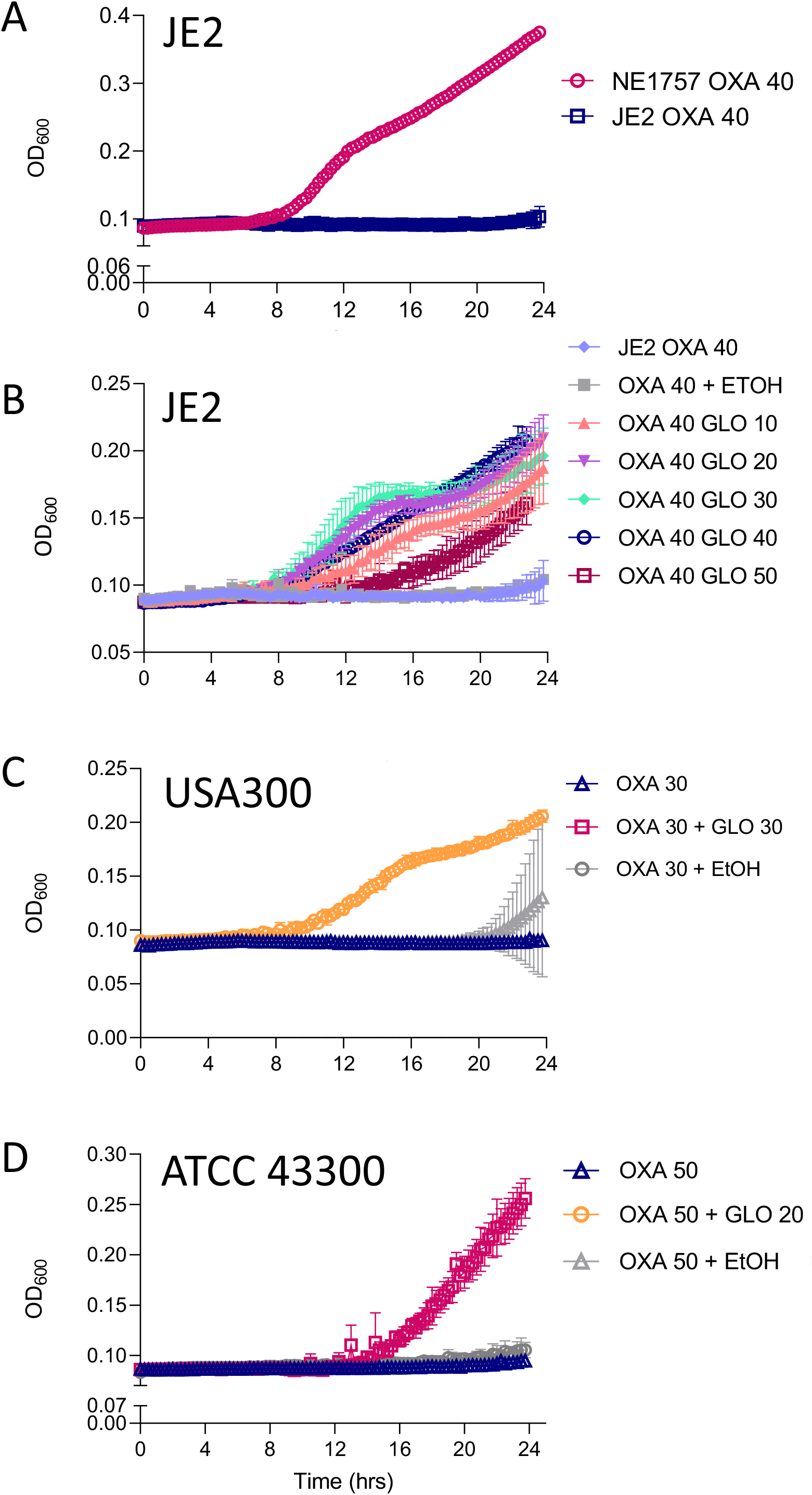
Mutation of *lspA* or exposure to globomycin increases oxacillin resistance. **(A)** JE2 and NE1757 (*lspA*) grown in MHB 2% NaCl supplemented with oxacillin 40 μg/ml. **(B)** JE2 grown in MHB 2% NaCl supplemented with oxacillin 40 μg/ml and globomycin concentrations ranging from 10-50 μg/ml or MHB 2% NaCl supplemented with oxacillin 40 μg/ml and 0.6% ethanol (control solvent for globomycin). **(C)** USA300 grown in MHB 2% NaCl supplemented with oxacillin 50 μg/ml, MHB 2% NaCl supplemented with oxacillin 50 μg/ml and globomycin 20 μg/ml or MHB 2% NaCl supplemented with oxacillin 50 μg/ml and 0.6% ethanol (control solvent for globomycin). **(D)** ATCC 43300 grown in MHB 2% NaCl supplemented with oxacillin 30 μg/ml, MHB 2% NaCl supplemented with oxacillin 30 μg/ml and globomycin 30 μg/ml or MHB 2% NaCl supplemented with oxacillin 50 μg/ml and 0.6% ethanol (control solvent for globomycin). The oxacillin and globomycin concentrations used in these experiments were determined empirically for each strain. Cultures were grown in a Tecan Sunrise incubated microplate reader for 24 h at 35°C. OD_600_ was recorded at 15 min intervals and growth curves were plotted in Prism software (GraphPad). The data presented are the average of 3 independent biological replicates, and error bars represent standard deviations.

These experiments were extended to USA300_FPR3757, from which JE2 is derived and ATCC43300, a *SCCmec* type II MRSA clinical isolate. Oxacillin concentrations of 50 μg/ml and 30 μg/ml inhibited growth of USA300 and ATCC4330, respectively (Fig. 5). Globomycin concentrations (determined empirically for each strain) of 20 μg/ml for USA300 (Fig. 4C) and 30 μg/ml for ATCC43300 (Fig. 4D) rescued growth in the presence of oxacillin. These data demonstrate that the increased oxacillin resistance phenotype observed in the *lspA* mutant NE1757, can be replicated by globomycin-induced inhibition of LspA activity in wild type JE2 and other MRSA strains.

**Fig. 5.**
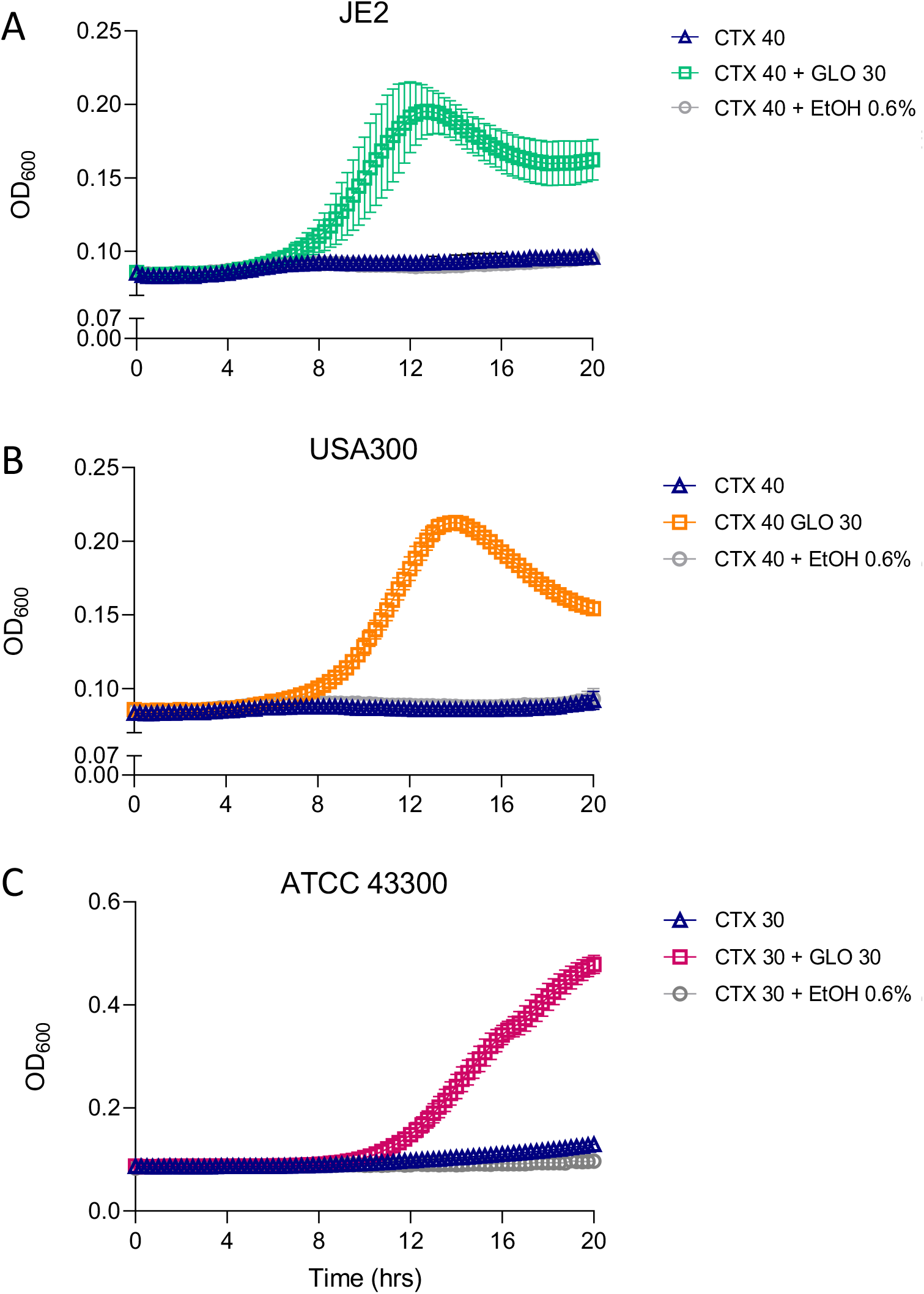
Globomycin increases cefotaxime resistance in JE2, USA300 and ATCC43300. **(A)** JE2 was grown in MHB 2% NaCl supplemented with cefotaxime (CTX) 40 μg/ml, MHB 2% NaCl supplemented with CTX 40 μg/ml and globomycin 30 μg/ml or MHB 2% NaCl supplemented with CTX 40 μg/ml and 0.6% ethanol (control solvent for globomycin). **(B)** USA300 was grown in MHB 2% NaCl supplemented with CTX 40 μg/ml, MHB 2% NaCl supplemented with CTX 40 μg/ml and globomycin 30 μg/ml or MHB 2% NaCl supplemented with CTX 40 μg/ml and 0.6% ethanol (control solvent for globomycin). **(C)** ATCC 43300 was grown in MHB 2% NaCl supplemented with CTX 30 μg/ml, MHB 2% NaCl supplemented with CTX 30 μg/ml and globomycin 30 μg/ml or MHB 2% NaCl supplemented with CTX 30 μg/ml and 0.6% ethanol (control solvent for globomycin). The cefotaxime and globomycin concentrations used in these experiments were determined empirically for each strain. The cultures were grown in a Tecan Sunrise incubated microplate reader for 24 h at 35°C OD_600_ was recorded at 15 min intervals and growth curves were plotted in Prism software (GraphPad). The data presented are the average of 3 independent biological replicates, and error bars represent standard deviations.

In contrast to the observation that globomycin increased β-lactam resistance in wild type JE2, USA300 and ATCC43300, the growth of NE1757 in a range of globomycin concentrations from 10 - 50 μg/ml had a dose-dependent and negative effect on growth in the presence of oxacillin 40μg/ml (Fig. S7). Taken together, these data suggest that globomycin antagonises β-lactam antibiotics, increasing MRSA resistance to oxacillin and cefotaxime in a LspA-dependent manner. However, in the absence of *lspA*, the combination of globomycin and oxacillin interferes with growth of JE2, particularly at higher concentrations of globomycin, perhaps due to off-target effects.

To determine if globomycin could also increase resistance to other classes of β-lactam antibiotics, its effect on cefotaxime resistance in JE2, USA300 and ATCC43300 was evaluated. Cefotaxime was chosen because it is a 3^rd^ generation cephalosporin with broad spectrum activity against Gram-positive and Gram-negative bacteria commonly used in the clinic, whereas oxacillin is a narrow-spectrum penicillin antibiotic. Cefotaxime 40 μg/ml inhibited growth of JE2 and USA300 (Fig. 5A and B), while 30 μg/ml inhibited growth of ATCC43300 (Fig. 5C). Globomycin (30 μg/ml) rescued growth of all three strains in oxacillin (Fig. 5A, B and C) suggesting that globomycin-mediated inhibition of LspA may antagonise the activity of many or all β-lactam antibiotics.

### Mutation of *lgt* increases susceptibility to the lipoteichoic acid synthase inhibitor Congo red

Congo red was recently identified as a selective inhibitor of lipoteichoic acid synthase (LtaS) activity (46). To investigate the possible involvement of LTA expression or stability in the increased resistance of the *lspA* mutant to β-lactams serial dilutions of overnight *lspA, lgt* and *lspA/lgt* mutant cultures were spotted onto TSA 0.125% Congo Red. Our data showed that mutation of *lgt* dramatically increased susceptibility to the selective LtaS inhibitor Congo red (Fig. 6) suggesting that impaired lipoprotein processing affects the expression or stability of LTA. However, while the *lgt/lspA* double mutant was even more susceptible to Congo red than the single *lgt* mutant, the single *lspA* mutations in NE1757 and NE1757 MM did not significantly alter Congo red susceptibility and complementation of NE1757 was also associated with increased susceptibility (Fig. 6). These data show that mutation of *lgt* or multicopy expression of *lspA* both increase susceptibility to this LTA inhibitor indicating that the lipoprotein processing pathway impacts LTA synthesis/stability in a complex manner. The mutations in the *lgt* and *lspA* genes alone, and in particular when combined, significantly increased resistance to the alanylation pathway inhibitor D-cycloserine (DCS)(Table 1). However lack of correlation between the effects of *lgt* and *lspA* mutations on susceptibility to Congo red, DCS and β-lactams indicates that further analysis is needed to better understand the interactions between lipoprotein processing pathway intermediates and the expression/stability of LTA and WTA.

**Fig. 6.**
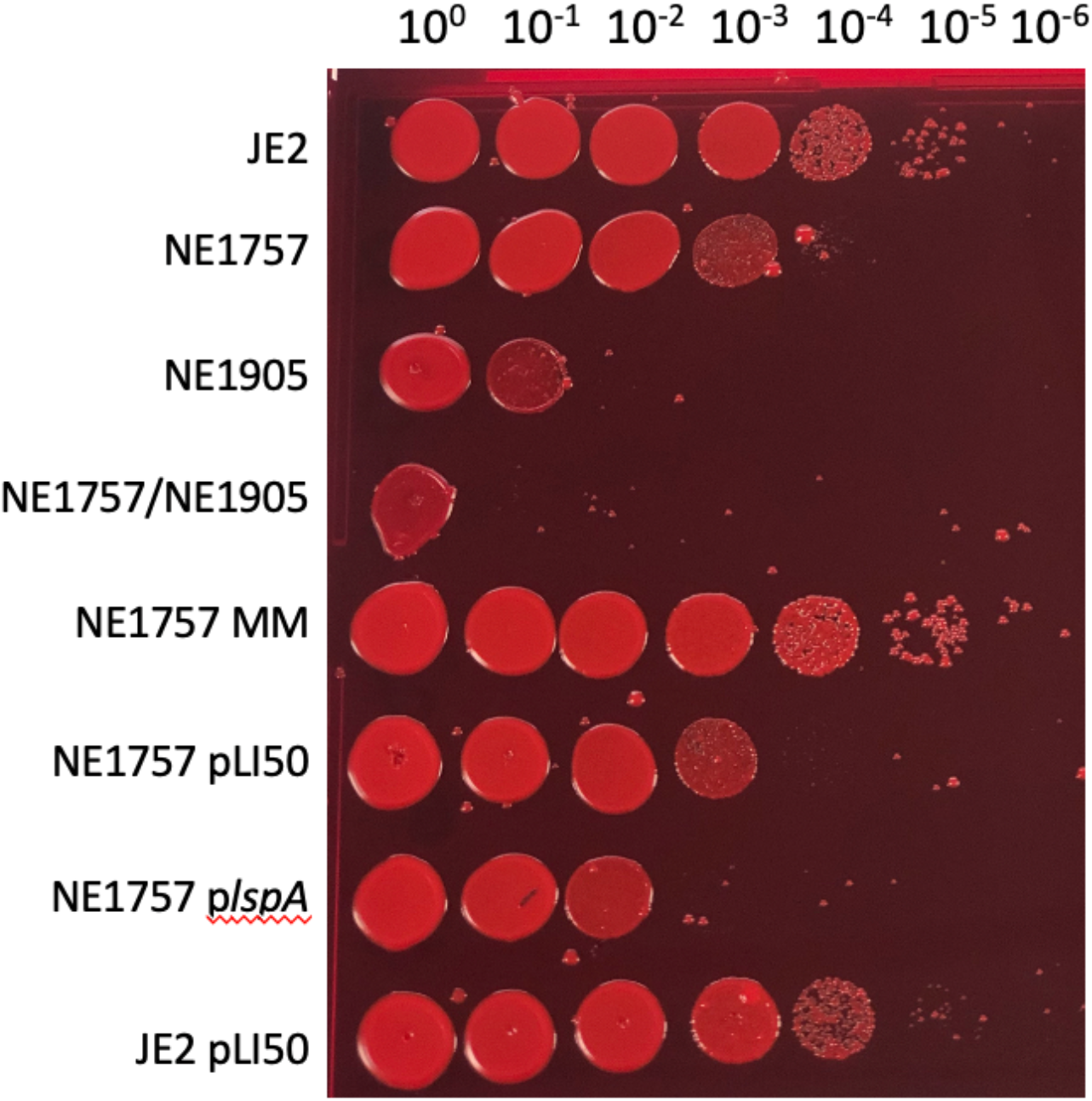
Mutation of *lgt* increases susceptibility to Congo red. Overnight cultures of JE2, NE1757 (*lspA*), NE1905 (*lgt*), NE1757/NE1905 *(lgt/lspA)*, NE1757 MM (markerless *lspA* mutation), NE1757 pLI50, NE1757 p*lspA* and JE2 pLI50 were grown in TSB (without antibiotic selection), normalised to an OD_600_ of 1 in PBS, serially diluted and spotted onto TSA plates supplemented with 0.125% Congo red, and the plates were incubated for 24 h at 37°C. A representative image is shown.

### Mutation of *lgt* in the *lspA* background restores wild type levels of β-lactam resistance

LspA catalyses the second major step in the lipoprotein processing pathway (Fig. 2). To probe the contribution of lipoprotein processing to LspA-controlled oxacillin resistance, we compared the impact of *lgt, lspA* and *lgt/lspA* mutants on growth and resistance to oxacillin, as well as cefotaxime, nafcillin and vancomycin. Lgt catalyses the addition of a diacylglycerol moiety onto preprolipoproteins, from which the signal peptide is then cleaved by LspA (Fig. 2). To construct a *lspA/lgt* double mutant the erythromycin resistance marker of the *lspA::Tn* allele in NE1757 was first exchanged for a markerless transposon to generate a strain designated NE1757 MM into which the erythromycin-marked *lgt*::Tn allele from NE1905 was transduced. Comparison of growth of JE2, NE1757, NE1757 MM, NE1905 and the *lgt/lspA* double mutant NE1757 MM /NE1905 revealed no significant differences MHB, MHB with 2% NaCl or TSB (Fig. 7A, B, C). However, consistent with previous analysis of a *lgt* mutant (35), the *lspA, lgt* and in particular the *lgt/lspA* mutants exhibited impaired growth in CDM (Fig. 7D), suggesting that aberrant lipoprotein processing may affect nutrient acquisition under substrate-limiting conditions.

**Fig. 7.**
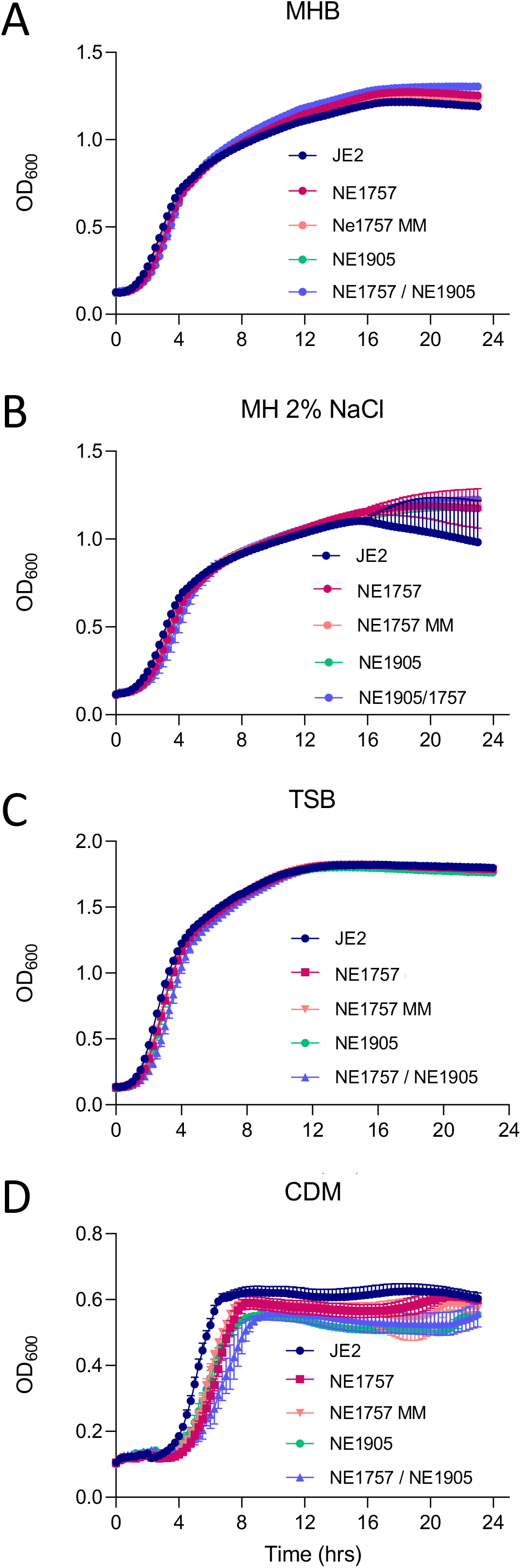
Mutation of *lspA* or *lgt* impacts growth in nutrient limited media but not in complex media. Growth of JE2 wild type, NE1757 (*lspA*::Tn), NE1757 MM (markerless *lspA*::Tn), NE1905 (*lgt*::Tn) and NE1757/NE1905 (markerless *lspA*::Tn / *lgt*::Tn) in Muller Hinton broth **(A)**, Muller Hinton broth with 2% NaCl **(B)**, TSB **(C)**, and chemically-defined medium **(D)**. Growth experiments were performed in 96-well hydrophobic plates in a Tecan Sunrise incubated microplate reader for 24 h at 37°C. OD_600_ was recorded at 15 min intervals and growth curves were plotted in Prism software (GraphPad). The data presented are the average of 3 independent biological replicates, and error bars represent standard deviations.

The *lgt* mutant NE1905 exhibited no changes in susceptibility to oxacillin, cefotaxime, nafcillin or vancomycin (Table 1). As observed for oxacillin, the *lspA* mutant NE1757 was more resistant to cefotaxime and nafcillin and these phenotypes were complemented by the *plspA* plasmid (Table 1). Neither the *lspA* nor *lgt* mutations increased resistance to vancomycin (Table 1). Oxacillin and nafcillin MICs were restored to wild type levels in the *lspA/lgt* double mutant, and the cefotaxime MIC was significantly reduced (Table 1).

Taken together, these data indicate that while mutation of *lspA* or *lgt* or both impact growth in CDM, only mutation of *lspA* alone is associated with increased β-lactam resistance. The *lgt* mutation and possible accumulation of unprocessed prolipoproteins (Fig. 8A,B) does not increase β-lactam resistance, whereas the possible accumulation of diacylglyceryl-lipoprotein in a *lspA* mutant (Fig. 8C, D) is associated with this phenotype. The oxacillin, cefotaxime and nafcillin MICs of the *ecsB* mutant from the NTML were the same as wild type (Table 1) indicating that downstream processing of the LspA-cleaved signal peptide is not associated with altered β-lactam resistance.

**Fig. 8.**
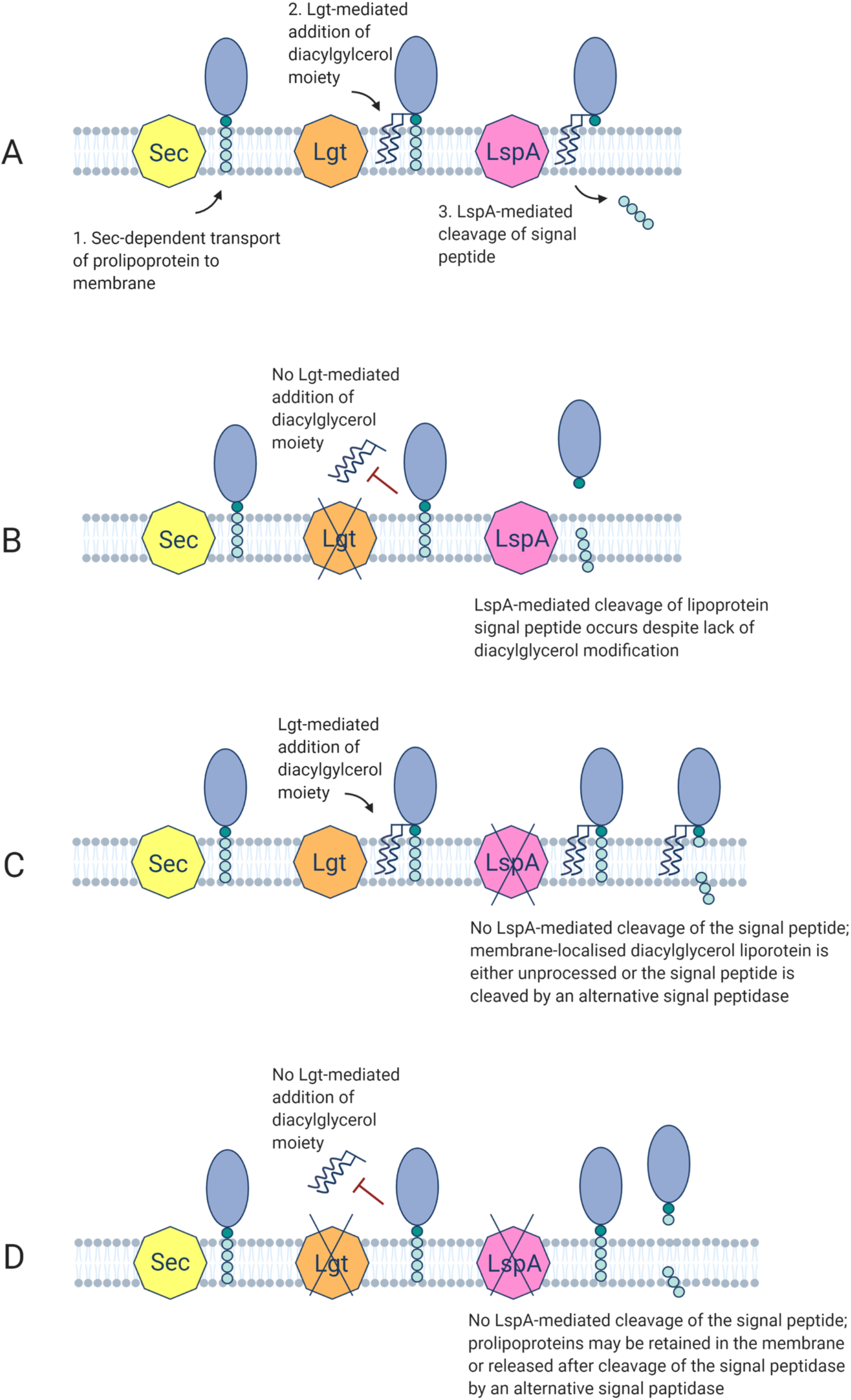
Predicted impacts of *lgt, lspA* and *lgt/lspA* mutations on lipoprotein processing in *S. aureus.* **(A)** Overview of lipoprotein processing in Gram-positive bacteria. **(B)** The *lgt* mutant lacks diacylglyceryl transferase activity and is unable to add the diacylglycerol molecule to the prolipoprotein, thus blocking accumulation of the LspA substrate. The prolipoprotein is likely to be released from the membrane due to LspA-mediated cleavage of the signal peptide. **(C)** LspA-mediated cleavage of the signal peptide is absent in the *lspA* mutant, but an alternative signal peptidase may undertake this activity. **(D)** Prolipoproteins are not processed in mutants lacking Lgt and LspA activity, and may be retained in the membrane or released after cleavage of the signal peptide by an alternative signal peptidase.

## Discussion

Advances in our understanding of the accessory factors that control levels of *mecA*/PBP2a-dependent resistance to methicillin has the potential to reveal new therapeutic targets and drugs that may facilitate the reintroduction of other β-lactam antibiotics for the treatment of MRSA infections. In this study, we demonstrated that mutation of the lipoprotein processing pathway gene *lspA*, or inhibition of LspA with globomycin increased resistance to β-lactam antibiotics. Although numerous mutations impacting the stringent response (ppGpp) and c-di-AMP signalling are associated with the transition from a heterogeneous to homogeneous pattern of resistance and elevated PBP2a expression (2,3), our data show that the *lspA* mutation was not associated with a HoR phenotype, increased PBP2a expression, or altered NaCl tolerance (which is controlled by c-di-AMP) (4,5). On the other hand, changes in β-lactam resistance independent of altered PBP2a regulation has long been known (47–49), and several auxiliary factors known to influence β-lactam resistance in MRSA have been described (49–54). In addition to unchanged PBP2a expression, no evidence of peptidoglycan remodelling was observed in NE1757 after growth in the presence or absence of oxacillin, potentially implicating wall teichoic acid (WTA) or lipoteichoic acid (LTA) synthesis or stability in the *lspA* mutant phenotype. Inhibition of WTA synthesis was previously shown to decrease β-lactam resistance in a PBP2a-independent manner (50). Reduced LTA stability, as evidenced by Western blotting and increased susceptibility to the selective lipoteichoic acid synthase inhibitor Congo red, was recently correlated with a PBP2a-independent reduction in β-lactam resistance in auxiliary factor *auxA* and *auxB* mutants (49). Interestingly AuxA is structurally similar to SecDF (49) and may interact with Sec pathway and lipoprotein processing (Fig. 8). The *lgt* mutant was significantly more susceptible to Congo red than the *lspA* mutant and multicopy expression of *lspA* also increased Congo red susceptibility suggesting that although the lipoprotein processing pathway modulates LTA synthesis/stability, the relationship between these two pathways appears to be complex.

Consistent with previous studies of lipoprotein pathway processing mutants in *S. aureus* (35), *Listeria monocytogenes* (55) and *Streptococcus agalactiae* (56), our analysis also showed that the *lgt, lspA* and *lgt/lspA* double mutants all exhibited impaired growth in CDM but not in TSB or MHB indicating that the impact of lipoprotein processing pathway mutations on nutrient acquisition and growth under nutrient limiting conditions can be compensated in rich media. Mutation of the *lgt* gene from the lipoprotein processing pathway (Figs. 2, 7) did not affect β-lactam resistance and introduction of the *lgt* mutation into a *lspA* mutant restored wild type levels of resistance. These data implicate accumulation of diacylglycerol lipoprotein in elevated β-lactam resistance. Consistent with this possibility, lipoproteins were retained in the membrane of a *S. agalactiae lspA* mutant, but were released into the supernatant in large concentrations by *lgt* and *lgt/lspA* mutants (56). Lipoproteins synthesised by the *S. agalactiae lspA* mutant retained their signal peptide, which was absent in a *lgt* mutant with LspA activity. Importantly, signal peptide processing also occurred in the *lgt/lspA* double mutant, albeit with cleavage occurring between different amino acids, implicating the involvement of an alternative signal peptidase. Our analysis revealed no change in oxacillin susceptibility in the MRSA *lgt/lspA* double mutant, indicating that even if an alternative peptidase can cleave the signal peptide, this may not impact β-lactam resistance. Furthermore, mutation of *ecsB*, which has recently been implicated in export of linear peptides from the lipoprotein processing pathway in *S. aureus* (32), did not change the oxacillin MIC in JE2 (Table 1) also indicating that downstream processing of signal peptides cleaved from lipoproteins by LspA is not associated with altered β-lactam resistance. In *L. monocytogenes*, deletion of *lgt* also led to significant release of lipoproteins into the supernatant. However, treatment of the *L. monocytogenes lgt* mutant with globomycin (inhibiting LspA activity) resulted in enhanced lipoprotein retention in the membrane (55), suggesting that the impact of globomycin and *lspA* mutation on lipoprotein processing is not necessarily the same. In a *S. aureus lgt* mutant, the Götz group reported that the majority of lipoprotein (lacking signal peptide) was released into the supernatant (35). Taken together, the data suggest that accumulation of membrane-anchored diacylglycerol lipoprotein with uncleaved signal peptide, or lipoprotein that is mis localised or released due to aberrant signal peptide processing by an alternative peptidase, is accompanied by increased β-lactam resistance in MRSA.

### Experimental procedures

#### Bacterial strains and culture conditions

Bacterial strains and plasmids used in this study are listed in Table S2. *Escherichia coli* strains were grown in Luria-Bertani (LB) broth or agar (LBA). *S. aureus* strains were grown in Tryptic Soy Broth (TSB), Tryptic Soy Agar (TSA) or chemically defined medium (CDM) (57). Muller Hinton Broth (MHB) or Muller Hinton Agar (MHA) (Oxoid) supplemented with 2% NaCl where indicated, were used for antimicrobial susceptibility testing (AST). Antibiotic concentrations used were 10 μg/ml erythromycin, 10 μg/ml chloramphenicol, 75 μg/ml kanamycin, 100 μg/ml ampicillin.

Two hundred and fifty ml flasks were filled with 25 ml growth media, and overnight cultures were used to inoculate the media at a starting OD_600_ of 0.05. Overnight cultures were grown in TSB, and washed once in 5 ml PBS before being used to inoculate the CDM cultures. Flasks were incubated at 37°C shaking at 200 rpm. OD_600_ readings were measured at 1 - 2 h intervals. Colony forming units (CFU) were enumerated in serially diluted 20 μl aliquots removed from flask cultures. Three independent biological replicates were performed for each strain and the resulting data plotted using GraphPad Prism software.

Data from growth experiments in a Tecan Sunrise microplate instrument were recorded by Magellan software. Overnight cultures were adjusted to an OD_600_ of 1 in fresh media and 10 μl inoculated into 200 μl media per well before being incubated at 37°C for 24 h with shaking. OD_600_ was recorded every 15 min. For CDM, overnight TSB cultures were first washed in 5 ml PBS and adjusted to OD_600_ of 1 in PBS.

#### Genetic manipulation of *S. aureus*

Phage 80α transduction was used to verify the association between antibiotic resistance phenotypes and transposon insertion-marked mutations from the NTML as described previously (39). Transductants were verified by PCR amplification of the target locus using primers listed in Table S3. The plasmid pTnT, which contains a truncated, markerless transposon was used to construct a markerless *lspA* mutant designated NE1757 MM, as described previously (58). A double *lgt/lspA* double mutant was subsequently constructed using phage 80α to transduce the *lgt*::Tn allele from NE1905 into NE1757 MM. A 1324 bp fragment encompassing the *lspA* gene was PCR amplified from JE2 genomic DNA using primers NE1757_INF#3_Fwd and NE1757_INF#3_Rev (Table 2), purified with GenElute™ PCR Clean-Up Kit and cloned into the *E. coli - Staphylococcus* shuttle vector pLI50 digested with *Eco*RI (New England Biolabs) using In-Fusion^®^ HD Cloning Kit (Clontech) to generate *plspA.* Using electroporation*, plspA* was transformed sequentially into Stellar™ *(E. coli* HST08) (Clontech), *S. aureus* RN4220 and finally NE1757.

#### Disk diffusion susceptibility assays

Cefoxitin disk diffusion susceptibility assays were performed in accordance with CLSI guidelines. Briefly, isolates were grown at 37°C on MHA for 24 h and 5 - 6 colonies were resuspended in 0.85% saline to OD_600_ of 0.08 - 0.1 (0.5 McFarland; 1 × 10^8^ CFU/ml) and swabbed onto MHA plates with a uniform agar depth of 4 mm. A 30 μg cefoxitin disk (Oxoid) was applied, the plate incubated at 35°C for 16 - 18 h and the zone of inhibition diameter measured. Strains were classified as sensitive, intermediate, or resistant, according to CLSI criteria.

#### Minimum inhibitory concentration (MIC) measurements

For oxacillin M.I.C.Evaluators (Oxoid) several colonies from 24 h MHA plates were resuspended in 0.85% saline to OD_600_ of 0.08 - 0.1, (0.5 McFarland standard) and evenly swabbed onto MHA 2% NaCl (4 mm agar depth). An M.I.C.Evaluator strip was applied and the plate incubated at 35°C for 24 h. Three biological repeats were performed for each strain.

For broth microdilution MIC measurements using 96-well plates, each plate row was used to prepare two-fold dilutions of antibiotic in MHB, typically ranging from 256 - 0.5 μg/ml across 10 wells. For oxacillin and nafcillin MIC measurements, MHB 2% NaCl was used. Several colonies from 24 h MHA plates were resuspended in 0.85% saline to OD_600_ of 0.08 - 0.1 (0.5 McFarland standard), diluted 1:20 in 0.85% saline and 10 μl of this cell suspension used to inoculate each well (approximately 5 × 10^4^ CFU/well) in a final volume of 100 μl. The plates were incubated at 35°C for 16 - 20 h, or 24 h incubation for oxacillin and nafcillin. The MIC was the lowest concentration of antimicrobial agent that completely inhibited growth.

Freshly prepared MHA plates (with 2% NaCl when using oxacillin and nafcillin) were supplemented with antimicrobial agents at 0.5, 1, 2, 4, 8, 16, 32, 64, 128 and 256 μg/ml to perform agar dilution MIC measurements. The inoculum was prepared as described for the broth microdilution method above before being further diluted 1:20 in 0.85% saline and 4 μl spot-inoculated onto each plate (approximately 10^4^ CFU per 5-8 mm diameter spot). The MIC was the lowest concentration of antimicrobial agent that completely inhibited growth after 16-18 h (24 h for oxacillin and nafcillin) at 35°C, disregarding a single colony or a faint haze associated with the inoculum.

Congo red susceptibility assays were performed as described previously (59). Briefly, overnight cultures grown in TSB were adjusted to an OD_600_ of 1 in PBS, serially diluted and spotted onto TSA plates supplemented with 1% aqueous Congo red solution (VWR) to a final concentration of 0.125%. Plates were incubated for 24 h at 37°C. These assays were performed with at least 3 biological repeats and representative image is shown for TSA 0.125% Congo red.

#### Autolytic activity assays

Overnight cultures (20 μl) were inoculated into 20 ml TSB, grown at 37°C (200 rpm) to OD_600_ of 0.5, washed with 20 ml cold PBS, resuspended in 1 ml cold PBS and finally adjusted to OD_600_ of 1. Triton X-100 was added at a final concentration of 0.1% v/v and the cell suspension incubated at 37°C with shaking (200 rpm). OD_600_ was recorded every 30 min for 4 h. Autolytic activity was expressed as a percentage of the initial OD_600_. NE406 (*atl*::Tn) was used as a control and at least 3 biological replicates performed for each strain.

#### Globomycin and β-lactam antibiotic synergy/antagonism assays

One hundred μl of MHB cefotaxime or MHB 2% NaCl oxacillin was added to the individual wells of 96-well plates. The oxacillin or cefotaxime concentration chosen for each strain was based on approximate MICs i.e. the lowest antibiotic concentration that inhibited growth. Overnight MHB cultures were resuspended in PBS at OD_600_ of 0.1 (0.5 McFarland standard) and then further diluted 1:20 before 10 μl (approximately 5 × 10^5^ CFU/ml) was added to each well and the plates were incubated at 35°C with shaking on a Tecan Sunrise microplate instrument for 20 h (cefotaxime) or 24 h (oxacillin). Globomycin ranging from 10 - 100 μg/ml was added to the cefotaxime or oxacillin cultures to measure potential synergism or antagonism. Three independent biological replicates were performed for each strain and antibiotic combination.

#### PBP2a western blot analysis

Overnight MHB cultures were used to inoculate 25 ml of MHB 2% NaCl, with or without 0.5 μg/ml oxacillin to a starting OD_600_ of 0.05, incubated at 35°C (200 rpm shaking) until an OD_600_ of 0.8 was reached before the cells were pelleted and resuspended in PBS to an OD_600_ of 10. Six μl of lysostaphin (10 μg/ml) and 1 μl of DNase (10 μg/ml) was added to 500 μl of this concentrated cell suspension before being incubated at 37°C for 40 min. Next, 50 μl of 10% SDS was added and the incubation continued for a further 20 min. The lysed cells were then pelleted in a microcentrifuge for 15 min, following which the protein-containing supernatant was collected and total protein concentration determined using the Pierce^®^ BCA Protein Assay Kit. Samples containing 8 μg total protein were mixed 1:1 with protein loading buffer (2x) (National Diagnostics) and incubated at 95°C for 5 min and loaded onto a 7.5% Tris-Glycine gel and separated at 120 V for 60 mins. Electrophoretic transfer to a PVDF membrane was carried out at 30 V for 30 min on the Trans-Blot Turbo Transfer System (Biorad). The PVDF membrane was blocked overnight in 5% skim milk powder in PBS at 4°C. The following day, the membrane was washed in fresh PBS. Anti-PBP2a (Abnova) was diluted 1:1000 in PBS-Tween 20 (0.1%) and incubated with the membrane for 1 h at room temperature. The membrane was washed in PBS to remove unbound antibody. The secondary antibody, HRP-rec-Protein G (Invitrogen) was diluted 1:2000 in PBS-Tween 20 (0.1%) and incubated with the membrane at room temperature for 1 h. Visualisation of the membrane was performed with the Opti-4CN Substrate kit (Biorad). Three independent experiments were performed and representative images of the developed PVDF membrane were recorded.

#### Population level antibiotic resistance profile analysis

Characterisation of the population resistance profile was performed as described previously (60). Overnight cultures were grown in TSB, adjusted to an OD_600_ of 1, 10-fold serially diluted from 10^-1^ to 10^-7^, and a 20 μl aliquot of each dilution plated onto a series of TSA agar plates supplemented with oxacillin 0.25, 0.5, 1, 2, 4, 8, 16, 32, 64 and 128 μg/ml. CFUs were enumerated after overnight incubation at 37°C and the results were expressed as CFU/ml at each oxacillin concentration. Three independent experiments were performed for each strain.

#### Antibiotic tolerance assay

Tolerance assays were performed as described previously (61). Briefly overnight TSB cultures were sub-cultured into 25 ml of fresh TSB in 250 ml flasks at a starting OD_600_ of 0.05 and grown to an OD_600_ of 0.5 at 37°C with 200 rpm shaking. At this time (T_0_) an aliquot was removed for CFU enumeration and 12.5 μg/ml oxacillin promptly added before the cultures were re-incubated. Antibiotic tolerance was expressed as the % CFU/ml after 2, 4, 6, 8 and 24 h growth in the antibiotic compared to the CFU/ml at T_0_. The results represent 3 biological replicates of each strain.

#### Salt tolerance assay

Overnight TSB cultures were adjusted to an OD_600_ of 1 in fresh TSB and serially diluted from 10^-1^ to 10^-7^. Four μl of each dilution was spot-inoculated onto TSA 2.2M NaCl plates and incubated at 37°C overnight.

#### Genomic DNA extraction and Whole Genome Sequencing (WGS)

Genomic DNA (gDNA) extractions were performed using the Wizard^®^ Genomic DNA Purification Kit (Promega) following pre-treatment of *S. aureus* cells with 10 μg/ml lysostaphin (Ambi Products LLC) at 37°C for 30 min. WGS was performed by MicrobesNG (http://www.microbesng.uk) using an Illumina sequencing platform with 2×250 bp paired-end reads. CLC Genomics Workbench software (Qiagen) was used for genome sequencing analysis of strains. As a reference genome, a contig was produced for wild type JE2 by mapping Illumina reads onto the closely related USA300 FPR3757 genome sequence (RefSeq accession number NC_07793.1). The Illumina short read sequences from NE1757 were then mapped onto the assembled JE2 sequence and the presence of the transposon insertion confirmed. Single Nucleotide Polymorphisms (SNPs), deletions or insertions were mapped in the NE1757 genome and presence of large deletions ruled out by manually searching for zero coverage regions using the CLC Genomics Workbench software.

#### Peptidoglycan analysis

For each strain and growth condition tested, independent quadruplicate 50 ml cultures were grown to an OD_600_ of 0.5, harvested and resuspended in 5 ml PBS before peptidoglycan was extracted as described previously (39,62). Mass spectrometry was performed on a Waters XevoG2-XS QTof mass spectrometer. Structural characterization of muropeptides was determined based on their MS data and MS/MS fragmentation pattern, matched with PG composition and structure reported previously (63–66).

## Acknowledgements

This study was funded by grants from the Health Research Board (HRA-POR-2015-1158 and ILP-POR-2019-102) (www.hrb.ie), the Irish Research Council (GOIPG/2016/36) (www.research.ie), Science Foundation Ireland (19/FFP/6441) (www.sfi.ie) to J.P.O’G, and Svenska Forskningsrådet Formas to F.C. We are grateful to Christopher Campbell and Laura Gallagher for assistance and advice throughout this study. The illustrations in Figures 2 and 8 were created with Biorender.com.

## Supplementary Tables and Figures

**Supplementary Table S1.** Genome sequence changes in NE1757 (*lspA*::Tn)

| Reference Position<sup>1</sup> | Type<sup>2</sup> | Ref.<sup>3</sup> | Allele<sup>4</sup> | Freq.<sup>5</sup> | Annotation |
| --- | --- | --- | --- | --- | --- |
| 1192002 - 1192003 | INS | TA | T-Tn-A | 100 | *Bursa aurealis* transposon |

<sup>1</sup>Reference position: Position in the USA300 FPR3757 genome sequence (NC\_007793.1).
<sup>2</sup>Type of mutation: INS, insertion.
<sup>3</sup>Ref: Nucleotide base in the USA300 FPR3757 reference genome.
<sup>4</sup>Allele: Nucleotide base at the same position in NE1757.
<sup>5</sup>Freq.: % Frequency at which INS was found in NE1757 compared to JE2

**Table S2.**
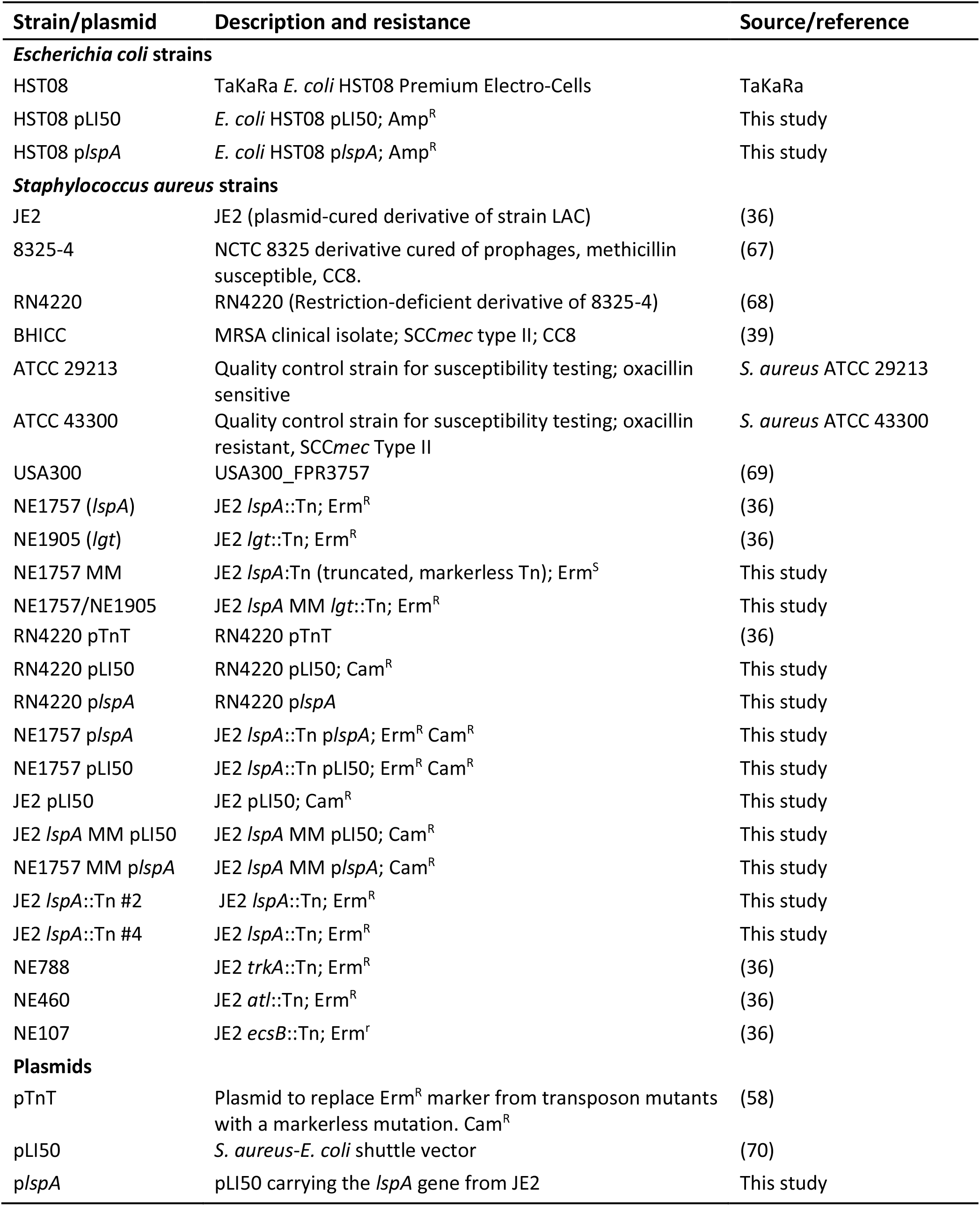
Bacterial strains and plasmids used in this study

| Strain/plasmid | Description and resistance | Source/reference |
| --- | --- | --- |
| ***Escherichia coli* strains** |  |  |
| HST08 | TaKaRa *E. coli* HST08 Premium Electro-Cells | TaKaRa |
| HST08 pLI50 | *E. coli* HST08 pLI50; Amp<sup>R</sup> | This study |
| HST08 p*lspA* | *E. coli* HST08 p*lspA*; Amp<sup>R</sup> | This study |
| ***Staphylococcus aureus* strains** |  |  |
| JE2 | JE2 (plasmid-cured derivative of strain LAC) | (36) |
| 8325-4 | NCTC 8325 derivative cured of prophages, methicillin susceptible, CC8. | (67) |
| RN4220 | RN4220 (Restriction-deficient derivative of 8325-4) | (68) |
| BHICC | MRSA clinical isolate; SCC*mec* type II; CC8 | (39) |
| ATCC 29213 | Quality control strain for susceptibility testing; oxacillin sensitive | *S. aureus* ATCC 29213 |
| ATCC 43300 | Quality control strain for susceptibility testing; oxacillin resistant, SCC*mec* Type II | *S. aureus* ATCC 43300 |
| USA300 | USA300_FPR3757 | (69) |
| NE1757 (*lspA*) | JE2 *lspA*::Tn; Erm<sup>R</sup> | (36) |
| NE1905 (*lgt*) | JE2 *lgt*::Tn; Erm<sup>R</sup> | (36) |
| NE1757 MM | JE2 *lspA*:Tn (truncated, markerless Tn); Erm<sup>S</sup> | This study |
| NE1757/NE1905 | JE2 *lspA* MM *lgt*::Tn; Erm<sup>R</sup> | This study |
| RN4220 pTnT | RN4220 pTnT | (36) |
| RN4220 pLI50 | RN4220 pLI50; Cam<sup>R</sup> | This study |
| RN4220 p*lspA* | RN4220 p*lspA* | This study |
| NE1757 p*lspA* | JE2 *lspA*::Tn p*lspA*; Erm<sup>R</sup> Cam<sup>R</sup> | This study |
| NE1757 pLI50 | JE2 *lspA*::Tn pLI50; Erm<sup>R</sup> Cam<sup>R</sup> | This study |
| JE2 pLI50 | JE2 pLI50; Cam<sup>R</sup> | This study |
| JE2 *lspA* MM pLI50 | JE2 *lspA* MM pLI50; Cam<sup>R</sup> | This study |
| NE1757 MM p*lspA* | JE2 *lspA* MM p*lspA*; Cam<sup>R</sup> | This study |
| JE2 *lspA*::Tn #2 | JE2 *lspA*::Tn; Erm<sup>R</sup> | This study |
| JE2 *lspA*::Tn #4 | JE2 *lspA*::Tn; Erm<sup>R</sup> | This study |
| NE788 | JE2 *trkA*::Tn; Erm<sup>R</sup> | (36) |
| NE460 | JE2 *atl*::Tn; Erm<sup>R</sup> | (36) |
| NE107 | JE2 *ecsB*::Tn; Erm<sup>r</sup> | (36) |
| **Plasmids** |  |  |
| pTnT | Plasmid to replace Erm<sup>R</sup> marker from transposon mutants with a markerless mutation. Cam<sup>R</sup> | (58) |
| pLI50 | *S. aureus*-*E. coli* shuttle vector | (70) |
| p*lspA* | pLI50 carrying the *lspA* gene from JE2 | This study |

**Table S3.**
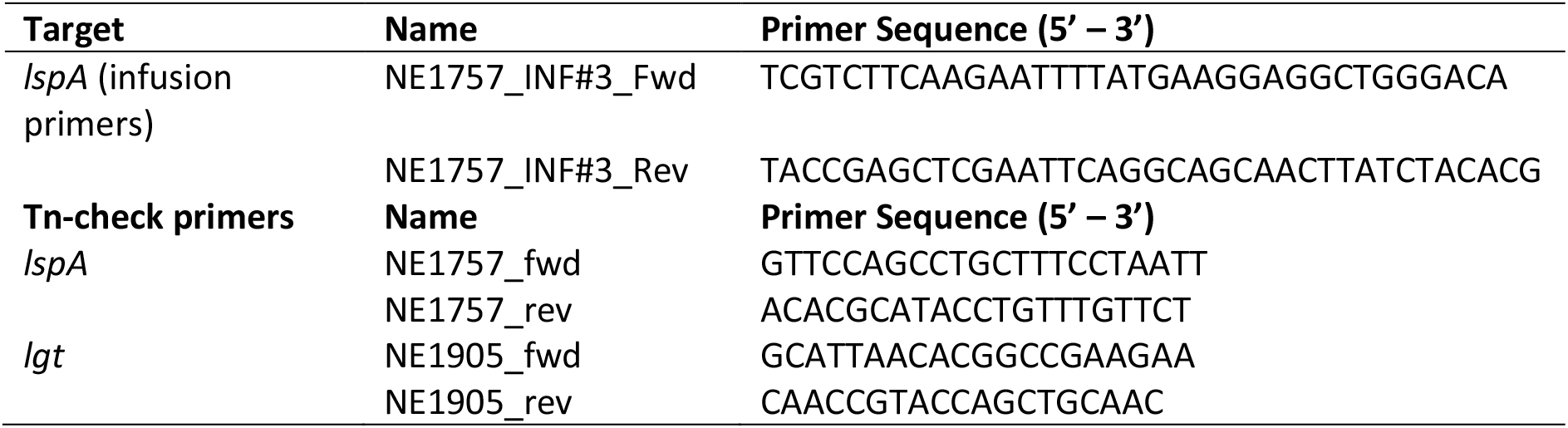
Oligonucleotides used in this study

**Supplementary Fig. S1.**
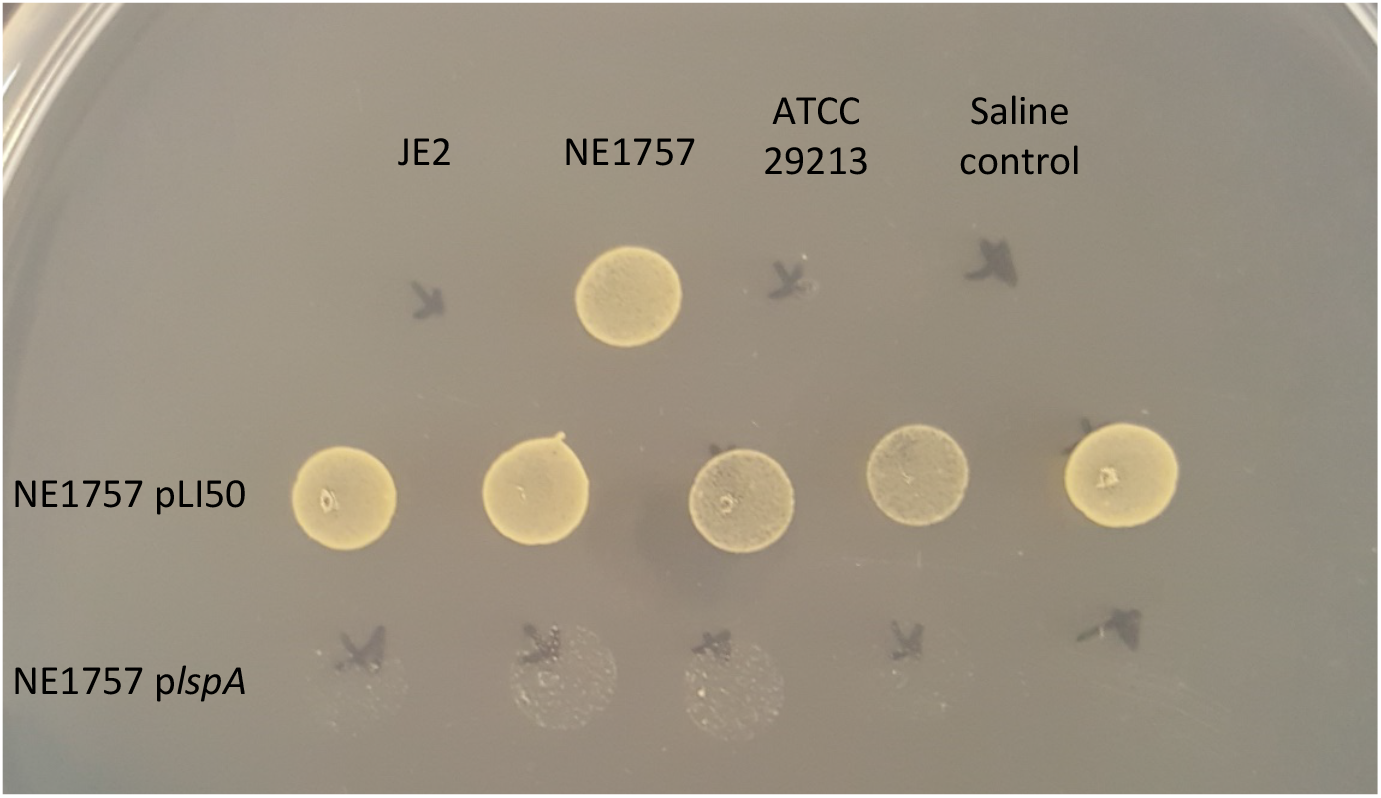
Complementation of the NE1757 mutant with the *lspA* gene is accompanied by a wild type oxacillin resistance phenotype. JE2, NE1757 (*lspA*::Tn), NE1757 pLI50 and NE1757 *plspA* were grown on MHA 2% NaCl supplemented with oxacillin 32 μg/ml. The plate shows 5 biological replicates each for NE1757 pLI50 and NE1757 *plspA.* ATCC 29213 was included as an oxacillin susceptible control.

**Supplementary Fig. S2.**
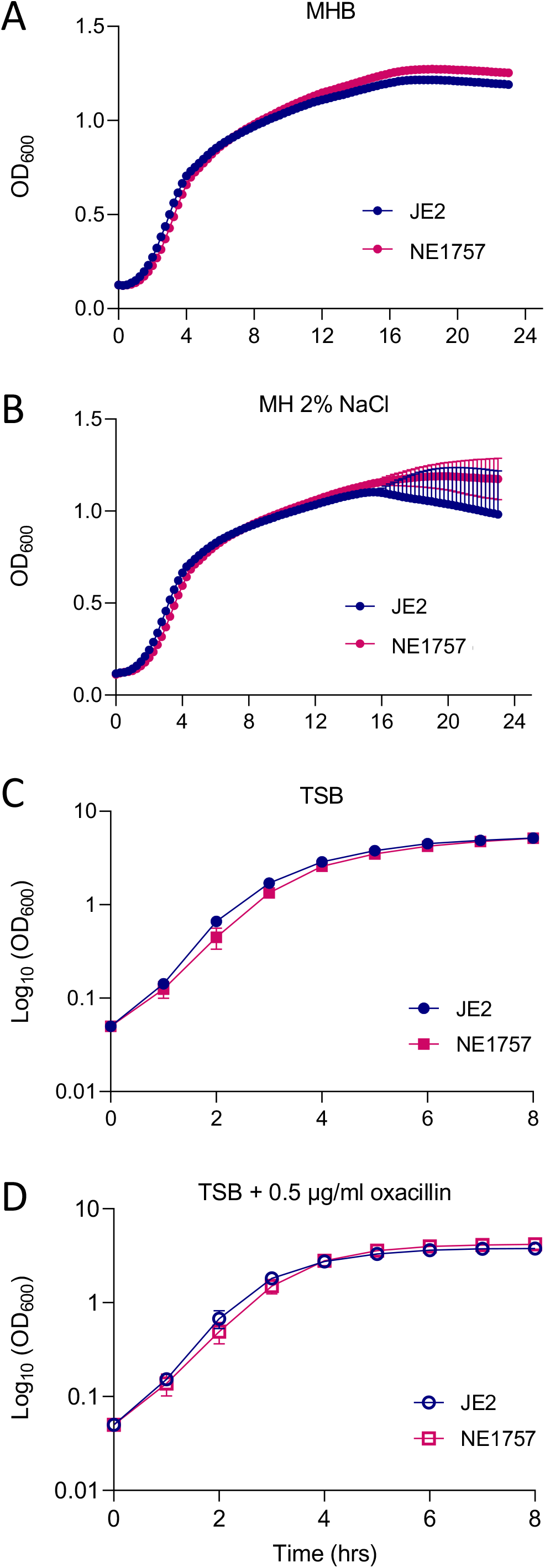
Mutation of *lspA* does not impact growth in MHB or TSB media. Growth of JE2 and NE1757 cultures in **(A)** MHB, **(B)** MHB 2% NaCl, **(C)** TSB and **(D)** TSB 0.5mg/ml oxacillin. MHB cultures were grown in 96-well plates in a Tecan Sunrise incubated microplate reader for 24 h at 37°C. The OD_600_ was recorded at 15 min intervals and growth curves were plotted in Prism software (GraphPad). TSB cultures were grown in flasks and the OD_600_ monitored every 2 h. All data presented are the average of 3 independent biological replicates, and error bars represent standard deviations.

**Supplementary Fig. S3.**
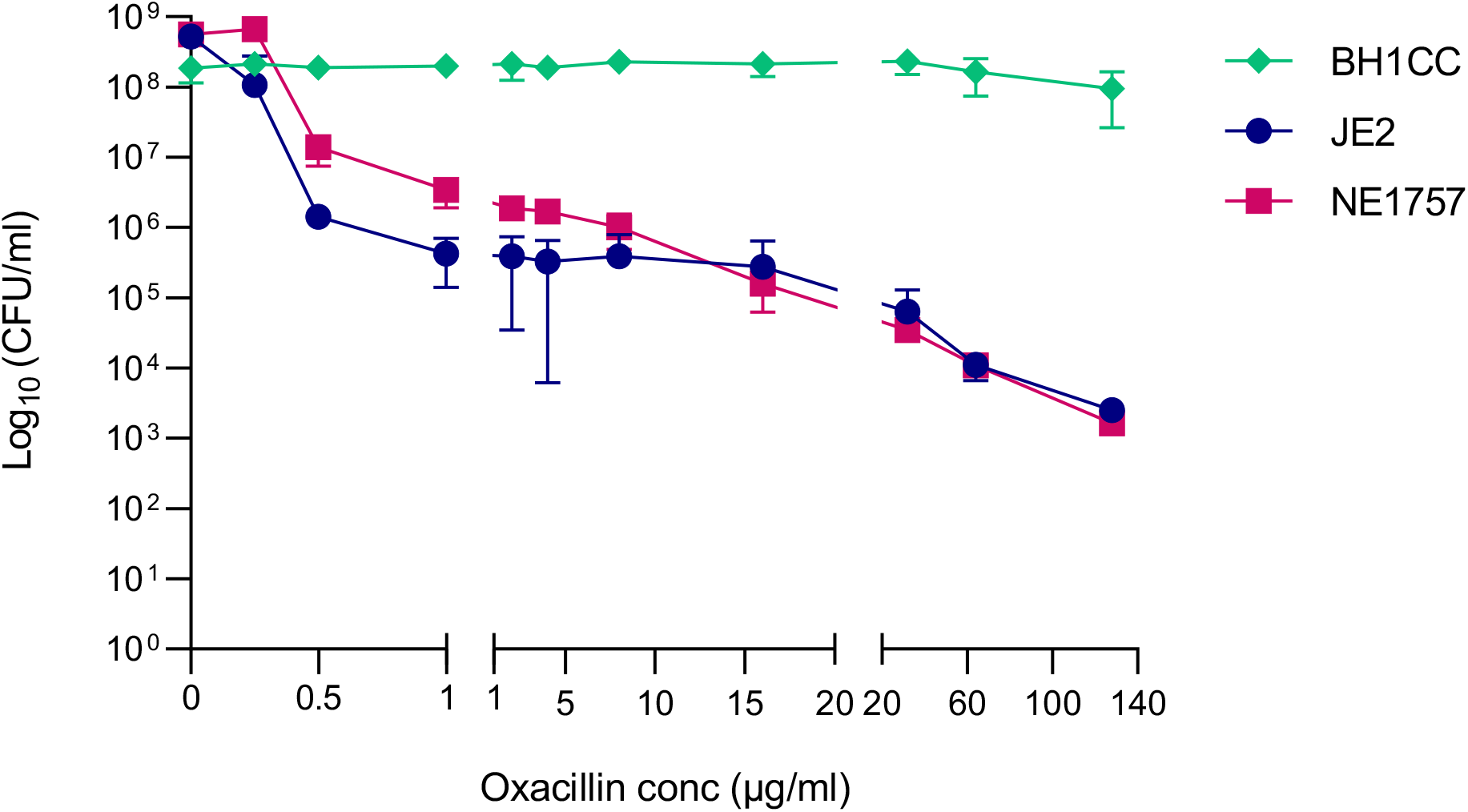
Population analysis profiling shows that the *lspA* mutant NE1757 exhibits heterogenous resistance to oxacillin. Overnight cultures of JE2, NE1757 and BH1CC were grown in TSB, adjusted to OD_600_ of 1, serially diluted and plated onto TSA and TSA supplemented with 0.25, 0.5, 1, 2, 4, 8, 16, 32, 64 and 128 μg/ml oxacillin. CFUs were enumerated after overnight incubation at 37°C. The data are expressed as CFU/ml at each oxacillin concentration, plotted using Prism software (GraphPad). Three independent experiments were performed, and error bars represent standard deviations. BH1CC, which exhibits homogenous oxacillin resistance, was included as a positive control.

**Supplementary Fig. S4.**
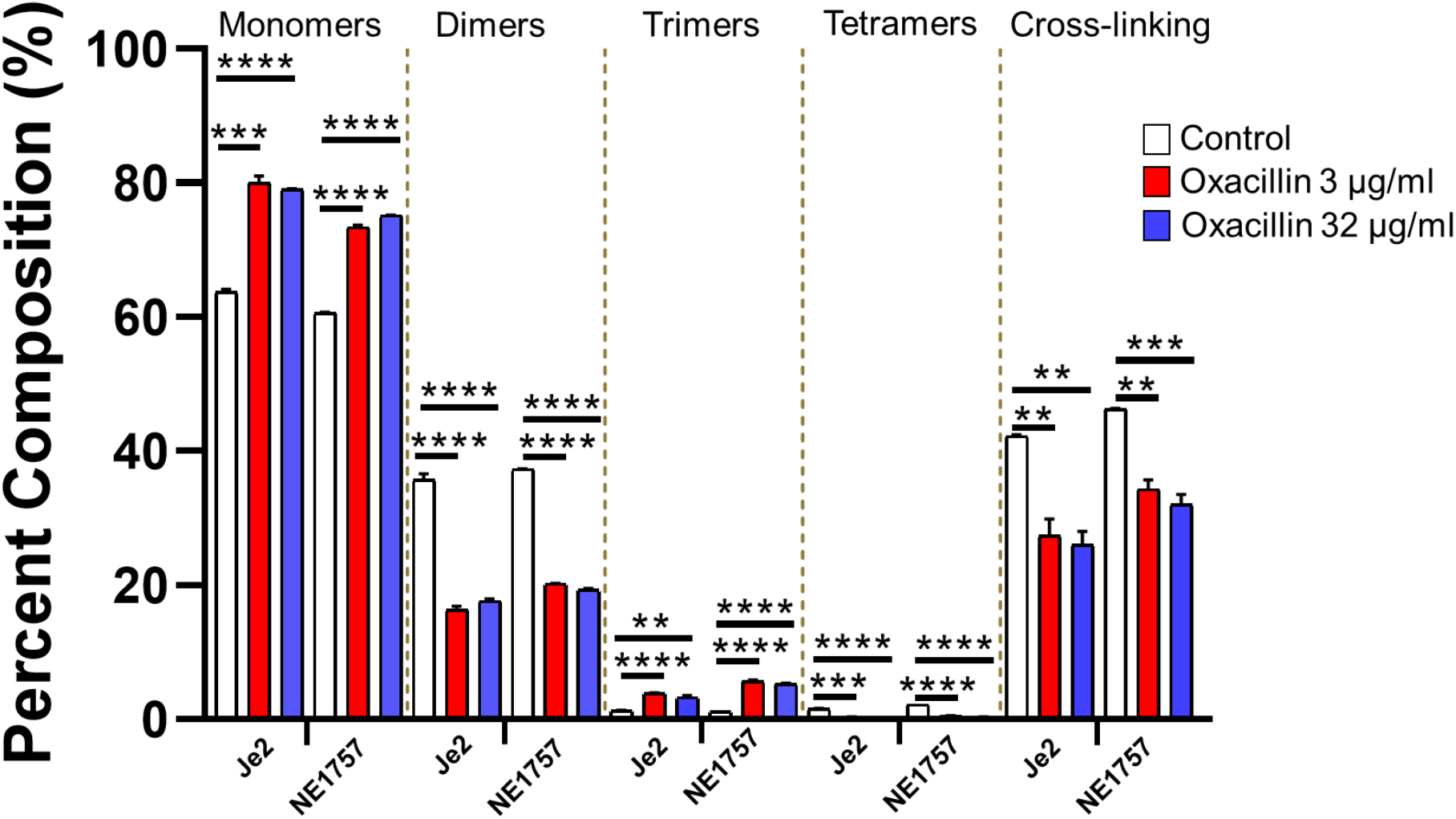
Relative proportions of cell wall muropeptide fractions based on oligomerization and relative cross-linking efficiency of cell wall muropeptide fractions in peptidoglycan extracted from JE2 and NE1757. Cells were collected from cultures grown to exponential phase in MHB or MHB supplemented with oxacillin 3 μg/ml or 32 μg/ml. Each profile shown is a representative of 3 biological replicates. Significant differences determined using Students t-test (**P < .01; ***P < .001; ****P<.0001).

**Supplementary Fig. S5.**
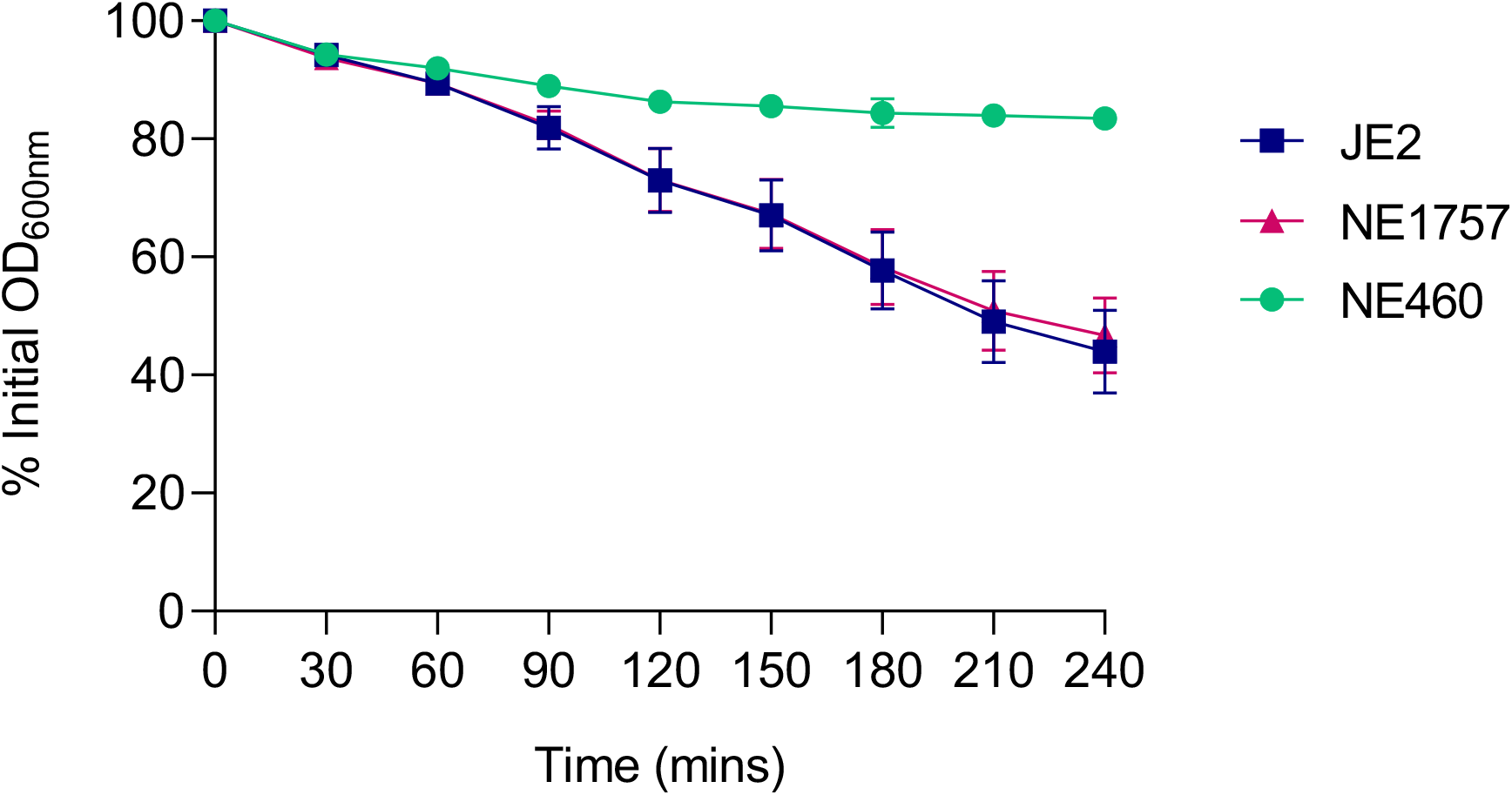
Autolytic activity is unaffected by the *lspA* mutation. Triton X-100-induced autolysis of JE2, NE1757 (*lspA::*Tn) and NE406 *(atl::Tn*, negative control). The strains were grown to OD_600_ = 0.5 in MHB medium at 37°C, before being washed in cold PBS and resuspended in 0.1% Triton X-100. The OD_600_ was monitored, and autolysis was expressed as a percentage of the initial OD_600_. The experiments were repeated 3 independent times, plotted using Prism software (GraphPad) and standard deviations are shown.

**Supplementary Fig. S6.**
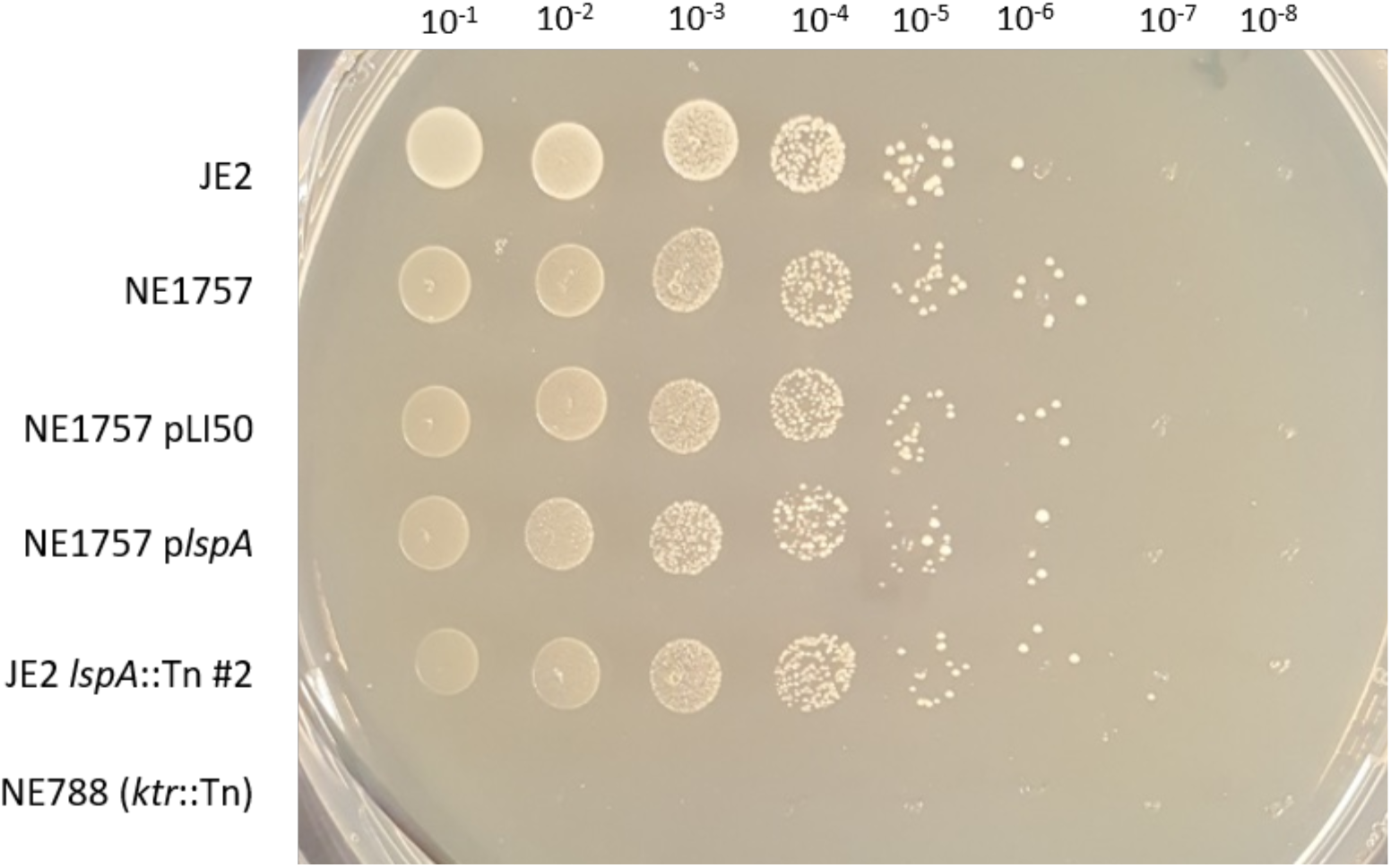
Mutation of *lspA* has no significant impact on salt tolerance in JE2. Overnight cultures of JE2, NE1757, NE1757 pLI50, NE1757 *plspA*, JE2 *lspA::Tn* #2 transductant and NE788 (JE2 *ktr::Tn*, NaCl-sensitive control) were grown in TSB and cell density was standardised to OD_600_ of 1. Four μl aliquots from 10-fold serial dilutions were spotted onto TSA supplemented with 2.2 M NaCl and the plates incubated overnight at 37°C. Three independent experiments were carried out and a representative image of a plate is shown

**Supplementary Fig. S7.**
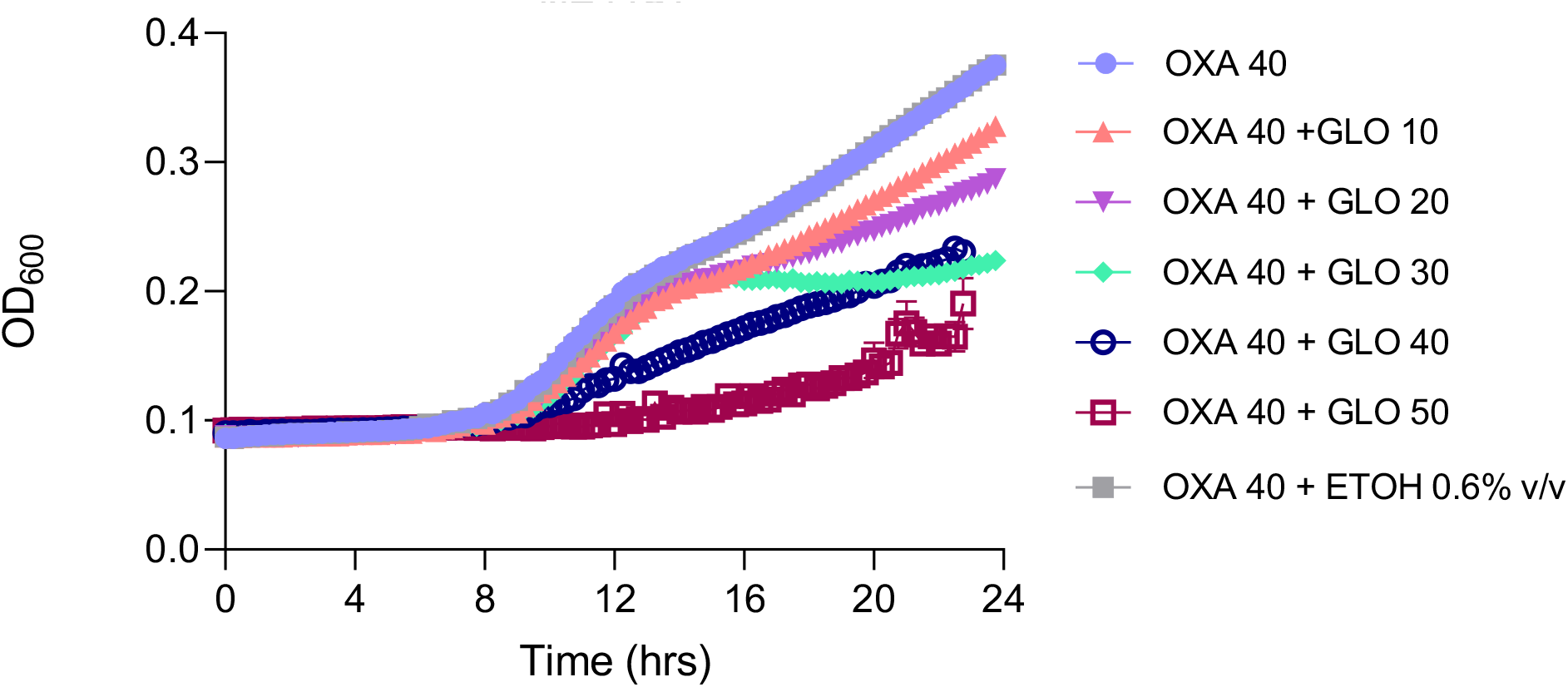
Globomycin does not increase oxacillin resistance in the *lspA* mutant NE1757. NE1757 was grown in MHB 2% NaCl supplemented with a sub-inhibitory concentration of oxacillin (40 μg/ml) and a range (10 - 50 μg/ml) of globomycin concentrations. The solvent for globomycin, 0.6% ethanol, was included as a control. The cultures were grown in a Tecan Sunrise incubated microplate reader for 24 h at 35°C. OD_600_ was recorded at 15 min intervals and growth curves were plotted in Prism software (GraphPad). The data presented are the average of 3 independent biological replicates, and error bars represent standard deviations.

